# Deep mutational scanning of CYP2C19 reveals a substrate specificity-abundance tradeoff

**DOI:** 10.1101/2023.10.06.561250

**Authors:** Gabriel E. Boyle, Katherine Sitko, Jared G. Galloway, Hugh K. Haddox, Aisha Haley Bianchi, Ajeya Dixon, Raine E. S. Thomson, Riddhiman K. Garge, Allan E. Rettie, Alan Rubin, Renee C. Geck, Elizabeth M. J. Gillam, William S. DeWitt, Frederick A. Matsen, Douglas M. Fowler

## Abstract

Cytochrome P450s (CYPs) are a family of enzymes responsible for metabolizing nearly 80% of small molecule drugs. Variants in CYPs can substantially alter drug metabolism, which may result in improper dosing and severe adverse drug reactions. CYPs have low sequence conservation, making it difficult to anticipate whether variant effects measured in one CYP may extend to others based on sequence alone. Even closely related CYPs, like CYP2C9 and its closest homolog CYP2C19, have distinct phenotypic properties despite sharing 92% amino acid sequence identity. Thus, we used Variant Abundance by Massively Parallel sequencing (VAMP-seq) to measure the steady-state protein abundance, a proxy for protein stability, of 7,660 missense variants in CYP2C19 expressed in cultured human cells. Our results confirmed positions and structural features critical for CYP function and revealed how variants at positions conserved across all eukaryotic CYPs influence abundance. We jointly analyzed 4,670 variants whose abundance was measured in both CYP2C19 and CYP2C9, finding that the homologs have different variant abundances in substrate recognition sites within the hydrophobic core, and that substitutions in some regions reduced abundance in CYP2C19 but not CYP2C9. We also measured the abundance of all single and some multiple WT amino acid exchanges between CYP2C19 and CYP2C9. While most exchanges had no effect, substitutions in substrate recognition site 4 (SRS4) reduced abundance in CYP2C19. When nearby amino acids were exchanged in double and triple mutants, we found distinct interactions between the sites in CYP2C19 and CYP2C9, revealing a region that is partially responsible for the difference in thermodynamic stability between the two homologs. Since these positions are also important for determining substrate specificity, there may be an evolutionary tradeoff between stability and altered enzymatic function. Finally, we used our data to analyze 368 previously unannotated human variants, finding that 43% had decreased abundance. Thus, by comparing variant effects between two closely related and important human genes, we have uncovered regions underlying their functional differences and paved the way for a more complete understanding of one of the most versatile families of enzymes.

## Introduction

Nearly 20,000 cytochrome P450 (CYP) heme monooxygenases have been identified across all domains of life^1^. CYPs catalyze a wide range of reactions with a diverse set of substrates, making them some of the most versatile enzymes in existence^2,3^. The 57 human CYP genes are grouped into 18 families with 43 subfamilies^4^, highlighting their genetic heterogeneity even within a single species. Despite their genetic and functional diversity, key structural and topological features of CYPs are highly conserved^5,6^. However, the relationship between CYP genetic variation, structure, and function is far from fully elucidated. For example, within CYP family 2, subfamily C (CYP2C), *CYP2C19* (MIM: <u>124020</u>, <u>609535</u>) and *CYP2C9* (MIM: <u>601130</u>) are the most closely related subfamily members, sharing 92% amino acid sequence identity. Their protein structures have nearly identical organization, with the largest deviations between their Cα backbones in the substrate binding cavity being only ∼3 Å^7^. Yet, the two homologs are functionally distinct, with largely disparate sets of substrates^8,9^ and divergent membrane interactions^10^. Moreover, CYP2C19’s melting temperature is ∼11℃ higher than CYP2C9’s^11^. Thus, even between these close homolog CYPs, the 43 diverged positions drive large functional differences.

Understanding the functional impact of variants across CYPs is particularly important because ∼12 of the 57 human CYPs contribute to metabolizing 70-80% of currently prescribed drugs that are processed by enzymes for elimination. Of those 70-80% CYP-metabolized drugs, CYP2C19 and CYP2C9 account for 20-30%^12^. Genetic variation in CYPs can substantially alter individual drug response leading to adverse drug reactions (ADRs), which are among the leading causes of morbidity and mortality^13,14^, and cost an estimated $30.1 billion annually^15^. To provide clinicians guidance for treating individuals with CYP variants, the Clinical Pharmacogenetics Implementation Consortium (CPIC) categorizes CYP genes into star (*) allele haplotypes according to enzymatic function: normal function, decreased function, no function, and increased function^16,17^. Genetic testing and employment of CPIC guidelines can prevent many ADRs. For example, up to 30% of the population may have a CYP2C19 variant with reduced function^18^ which may result in impaired activation of the antiplatelet drug clopidogrel. Genotyping for CYP2C19 loss of function variants can avoid major adverse cardiovascular events^19–21^. However, only a very small number of CYP variants have established functional consequences, and it is unknown to what degree variant effects in one CYP can be applied to others.

Measuring CYP variant function individually is laborious and low throughput, but massively multiplexed methods can be used instead. Variant Abundance by Massively Parallel sequencing (VAMP-seq) measures steady-state protein abundance of thousands of variants in parallel^22^. In VAMP-seq, as in other similar methods^23–26^, steady-state protein abundance is used as a proxy for protein stability. Steady-state protein abundance refers to the final concentration of a protein when its rates of synthesis and decay are balanced^27^. To control for variation in protein synthesis, the VAMP-seq vector uses a single promoter to express two fluorescent reporters from a single mRNA transcript using an internal ribosome entry site (IRES). In each cell, the fluorescent signal of the target gene fused to enhanced green fluorescent protein (eGFP) is normalized to mCherry fluorescence meaning changes in fluorescence are due to degradation. The resulting measurements correlate with changes in thermodynamic stability, indicating that reduced steady-state abundance is likely due to loss of protein stability^22,24,28,29^.

Previously, we used the VAMP-seq assay^22^ to measure the abundance of 6,370 of 9,780 possible missense variants in CYP2C9^30^. From the resulting variant effect map, we identified patterns of loss of abundance that revealed mutationally sensitive regions of the protein. Additionally, we revealed hundreds of variants with reduced abundance in the human population in addition to providing variant effect measurements for thousands of variants not yet observed^30^.

Here, we used VAMP-seq to measure 7,660 of 9,780 possible missense variants of CYP2C19. We identified 4,698 variants that likely result in reduced protein abundance, with 1,122 of those exhibiting complete abundance loss equivalent to nonsense mutations. We first analyzed positions conserved across all eukaryotic CYPs, revealing that all but six of the 58 conserved positions were intolerant of substitutions. Four of the tolerant positions were catalytically important sites buried in the hydrophobic core where mutations are nearly always deleterious, suggesting that some sites critical for enzyme function may not impact abundance. We jointly analyzed the CYP2C19 and CYP2C9 variant abundance dataset, and found 2,366 variants whose abundance differed between the two enzymes. Most differences were of small effect, though a fraction of the differences were large. While nearly all sites had at least one variant that differed, 83 of 489 (17%) sites were significantly different between the homologs. CYP2C9 had higher mutational tolerance in its hydrophobic core than CYP2C19, and variants in the structurally conserved K’ helix are highly deleterious in CYP2C19, but tolerated in CYP2C9, even though all K’ positions contain the same amino acid in both homologs. We analyzed WT amino acid exchanges between CYP2C19 and CYP2C9, revealing that sequence differences in a set of diverged positions partially drive the differences in thermodynamic stability between the homologs. These divergent positions are also important for substrate specificity, suggesting that reduced thermodynamic stability in CYP2C9 may have been evolutionarily tolerated in exchange for functional benefit^31^. Finally, we analyzed the effects of human CYP2C19 variants. Our abundance scores are largely concordant with existing functional annotations indicating that, like for many other proteins, loss of abundance accounts for the majority of loss-of-function alleles. We provided abundance scores for 368 out of 408 (90.2%) previously unannotated missense variants in the gnomAD database. Thus, by conducting the first comparative analysis of closely related CYPs using large-scale variant effect data we provide fundamental insights into common CYP structural features that differentially impact abundance between CYP2C19 and CYP2C9. We also provide functional annotations for human CYP2C19 variants which could be used to improve genotype-guided dosing of drugs metabolized by CYP2C19.

## Results

### Multiplexed measurement of CYP2C19 variant abundance

We used VAMP-seq to simultaneously measure the steady-state abundance of CYP2C19 variants in cultured human cells^22,30^ (**Fig. 1A**). VAMP-seq relies on two fluorescent reporters: GFP fused to each CYP2C19 variant to read out abundance, and mCherry expressed via an internal ribosome entry site (IRES) as a transcriptional control. Because CYP2C19 is N-terminally inserted into the endoplasmic reticulum membrane, we fused GFP onto the C-terminus, as we did for a previous VAMP-seq experiment on CYP2C9^30^. Expression of the wild type (WT) CYP2C19 C-terminal GFP fusion led to strong fluorescent signal, and R433W, a known destabilizing CYP2C19 variant, had substantially lower signal indicating that the C-terminal GFP fusion construct was compatible with VAMP-seq (**Fig. 1B**).

**Figure 1.**
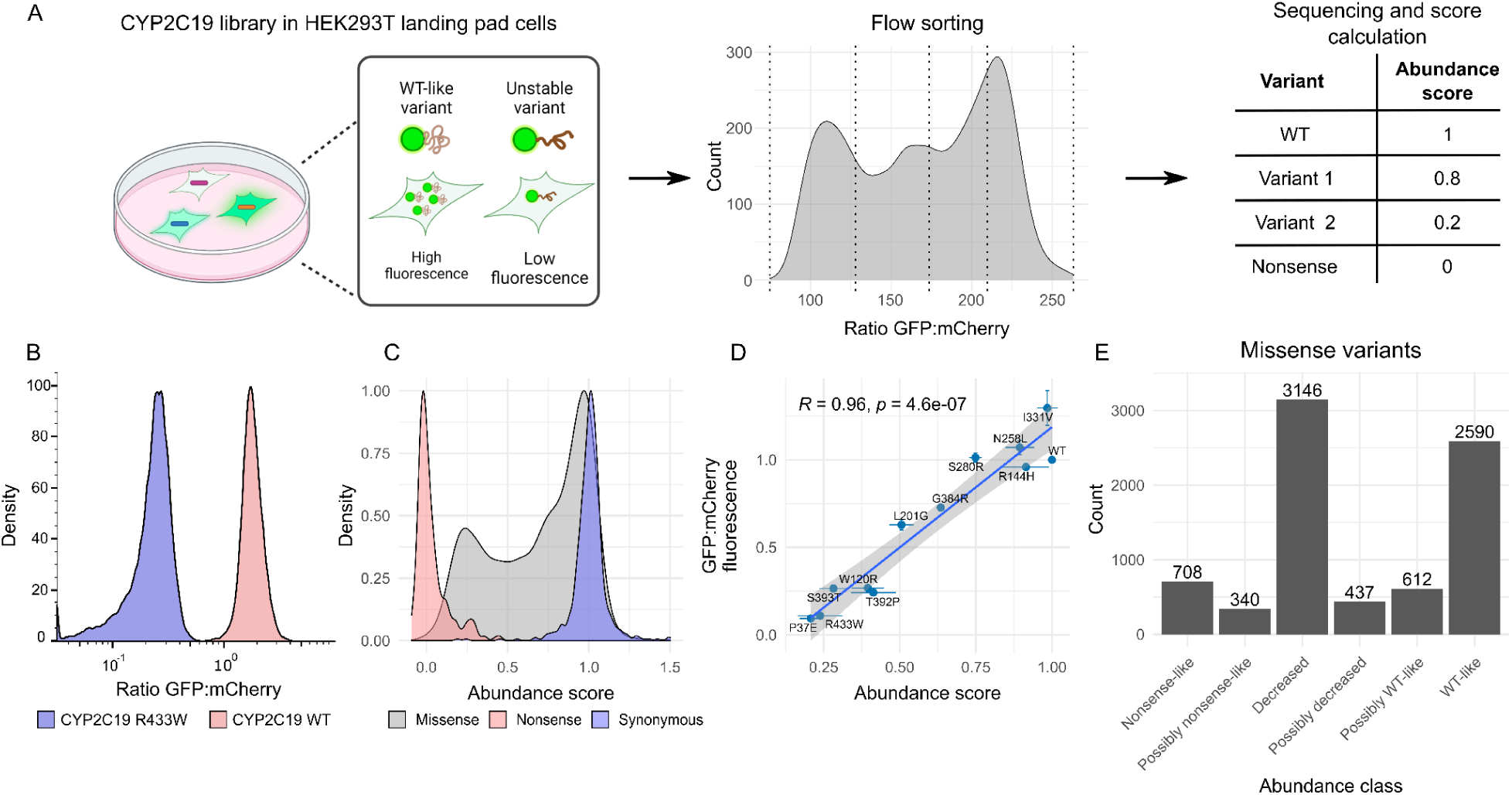
Multiplexed measurement of CYP2C19 abundance. Variant assessment by massively parallel sequencing (VAMP-seq) measures variant abundance at scale. A) In VAMP-seq, a barcoded library fused to GFP is recombined into a genomically integrated landing pad in HEK293T cells. mCherry is expressed co-transcriptionally via an IRES. Unstable variants are degraded by the proteostasis machinery of the cell, resulting in lower GFP signal compared to wild type (WT)-like variants. Flow cytometry is then used to sort cells into quartile bins according to fluorescence, bins are deeply sequenced, and barcode counts are used to calculate an abundance score. B) GFP:mCherry ratio for cells expressing either CYP2C19 WT (red) or the R433W destabilizing variant (blue) (n ∼30,000). C) Abundance score distributions for synonymous (n = 504), nonsense (n = 316) and missense (n = 7,660) variants. D) GFP:mCherry ratios, measured for cells using flow cytometry, for 10 individual variants plotted against their VAMP-seq-derived abundance scores (Pearson’s R=0.96, n = 30,000 cells). Error bars represent standard deviation of abundance scores (x-axis) or mean fluorescence (y-axis). E) Number of missense variants in each abundance class.

We introduced a barcoded library of CYP2C19 variants into HEK293T cells using a recombinase-based landing pad, such that each cell expressed only one variant^32,33^. Cells were sorted into quartile bins based on the ratio of GFP:mCherry fluorescence. Each bin was deeply sequenced, variant-associated barcodes were counted, and abundance scores were calculated based on weighted average of barcode frequencies across bins (**Fig. 1A**). Abundance scores were highly correlated between seven replicate sorting experiments arising from three independent library recombinations (**Supp.** Fig. 1A-D; Pearson’s *R* = 0.82 – 0.98). Replicate scores were averaged, filtered (**Supp.** Fig 2A-D) and normalized such that the median nonsense variant had a score of 0 and WT had a score of 1^22,30^.

Our final data set contained abundance scores for 8,480 of 10,290 (82%) possible variants, of which 7,660 were missense, 316 were nonsense, and 504 were synonymous. Abundance scores of synonymous and nonsense variants were well separated, with the missense variant distribution spread between nonsense and synonymous variants (**Fig. 1C**). Individually measured GFP:mCherry ratios for 10 variants spanning the range of abundance scores were highly correlated with VAMP-seq scores (**Fig. 1D**; Pearson’s R= 0.92). Our results are also highly consistent with a smaller scale VAMP-seq experiment encompassing 121 variants^34^ (**Supp.** Fig. 3A, Pearson’s R = 0.74). Thus, our VAMP-seq derived abundance scores faithfully reproduced variant abundance. Lastly, we classified variants according to their abundance score relative to the range of scores from nonsense and synonymous variants (**Supp.** Fig. 3B, **Fig. 1C,E**). The majority (58%, 4,620 variants) of missense variants decreased abundance (**Fig. 1E**).

### Mutational tolerance at conserved CYP2C19 positions reflects function

We visualized the abundance scores as a variant effect map (**Fig. 2A**) and projected position-averaged scores onto the CYP2C19 structure (**Fig. 2B**). Many of the low abundance variants occur within α-helices and β-sheets (**Fig. 2A-B**), especially in amino acids on interior α-helix turns and in regions closer to the protein core (**Fig. 2B**).

**Figure 2.**
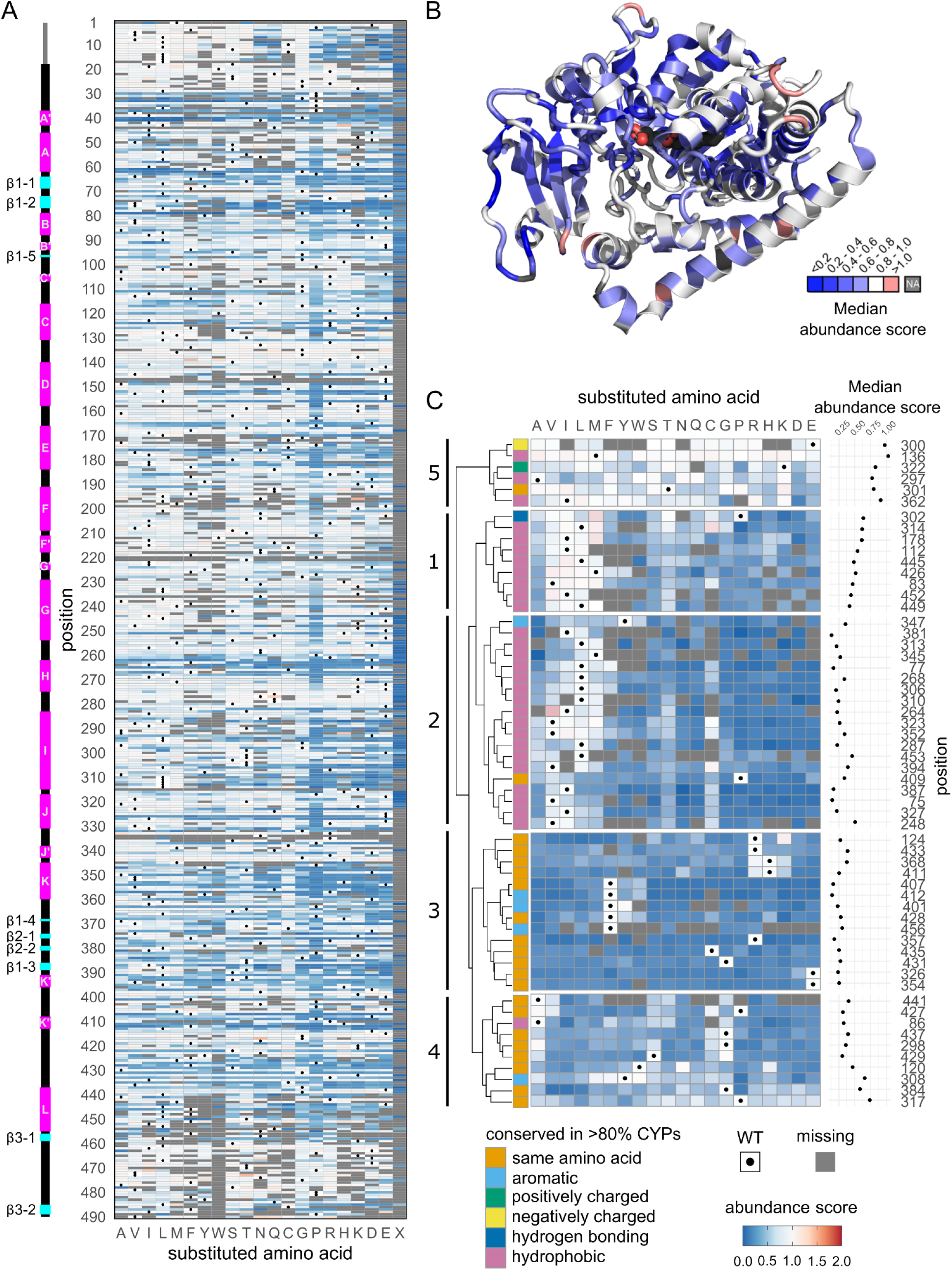
CYP2C19 variant abundance scores emphasize essential roles of conserved sites. A) Heatmap of CYP2C19 abundance scores. WT amino acids are represented by black dots, and missing data are shown in gray. Substituted amino acids are represented by their single letter abbreviations with “X” denoting a premature stop.Scores range from reduced abundance (blue) to increased (red). Secondary structure of CYP2C19 represented above the heatmap with α-helices shown in magenta and β-sheets shown in cyan. B) Median abundance scores for each position projected onto the CYP2C19 crystal structure (PDB: <u>4GQS</u>). Color represents binned median score, with missing scores represented in gray. Heme is colored by element (carbon: black, nitrogen: blue, oxygen: red, iron: yellow). C) Hierarchical clustering of CYP2C19 abundance score profiles by Euclidean distance at positions where >80% of eukaryotic CYPs had the same amino acid (orange) or >80% eukaryotic CYPs had amino acids with the same biophysical property (aromatic: light blue, positively charged: green, negatively charged: yellow, hydrogen bonding: dark blue, hydrophobic: pink)^38^. Cluster numbers labeled left of the dendrogram.

While CYPs vary widely in sequence, key structural and functional features are highly conserved^35,36^. However, despite this high level of conservation, the role of some positions in human CYPs are still poorly understood because some of these positions have not been studied, and others have only been studied in evolutionarily distant, non-human CYPs^37^. To bridge this gap, we investigated the abundance of variants at positions that are conserved across eukaryotic CYPs, defined as positions where >80% of CYPs have the same or biophysically similar amino acids^38^. Hierarchical clustering of these eukaryotically conserved positions revealed five clusters with distinct patterns of variant abundance scores (**Fig. 2C, Supp.** Fig. 4A). Overall, nearly all these conserved positions are critical for abundance. The clusters were defined by positions having similar variant effects amongst biophysically related amino acids^38^.

In clusters 1, 2, and 3 nearly all substitutions, except those of the same biophysical type, reduced abundance (**Fig. 2C, Supp.** Fig. 4A). In cluster 4, substitutions caused moderate loss of abundance, with no consistent pattern across all positions. The sole exceptions were two of the three positions where glycine was the WT amino acid. These positions tolerated alanine and cysteine substitutions suggesting that amino acid size is an important factor. Cluster 5 contained M136, A297, E300, T301, K322 and I362, all of which were substantially more tolerant of mutations than the other conserved sites indicating that they are critical to CYP2C19 function but not abundance. The combined conservation and tolerance of M136 and K322 can be explained by the fact that these positions are located on the surface of the protein and that they likely bind to the critical cofactor cytochrome P450 reductase (CPR), as they do in the closely related CYP2C9^39,40^. However, amino acids A297, E300, T301, and I362 are buried in the hydrophobic core making their mutational tolerance more challenging to explain (**Supp.** Fig 4B). Positions 297 and 362 influence substrate specificity, and >80% eukaryotic CYPs have hydrophobic amino acids at these positions^38^. Surprisingly, while substitutions are tolerated at these positions, some hydrophobic substitutions elicit moderate reductions in abundance. T301 is a critical threonine for oxygen activation and catalysis in 2C19^7,41–43^ and contains hydrogen-bonding amino acids in >80% eukaryotic CYPs at this position, and most substitutions did not appreciably reduce abundance. Finally, E300 stabilizes a water network during proton delivery^43^, and substitutions other than aspartic acid were tolerated at this position.

Thus, substitutions at nearly all conserved positions caused reduced abundance. However, positions 136, 297, 300, 301, 322, and 362, which participate in catalysis or cofactor binding, were largely tolerant of substitutions despite their location in the hydrophobic core of CYP2C19. We speculate that this tolerance is a consequence of the dynamic and flexible nature of CYP active sites, making these positions important for catalytic activity but not folding and stability^44^.

### Comparing variant abundance effects between CYP2C19 and CYP2C9 reveals core-stabilizing regions with distinct mutational tolerance

Next, we investigated variant effect patterns in CYP2C19 compared to its closest homolog, CYP2C9. CYP2C19 and CYP2C9 share 92% protein sequence identity and nearly identical crystal structures (**Supp.** Fig. 5, RMSD = 0.596 Å). However, they have important functional differences, notably their substrate profiles and membrane interactions^8,10,45,46^. Moreover, the temperature at which they lose the ability to bind their heme cofactor, which reflects thermodynamic stability^47^, differs by 11°C^11^. Thus, small differences in sequence and structure translate into distinct functional and phenotypic characteristics.

To understand how these functional differences arise, we sought to estimate how much the abundance score is shifted between the CYP2C19 abundance data presented here and abundance data from a previous VAMP-seq experiment we conducted on CYP2C9 variants^30^. The combined dataset contained 4,670 variants whose abundance was scored in both CYPs. Most variants had similar effects in both homologs, though a subset of variants had large shifts in effects (**Fig. 3A**, Pearson’s r = 0.77). While some shifts are likely due to actual biological differences, others are due to the noise inherent in any high-throughput experiment. To identify which shifts are most likely due to signal, we inferred shifts using a joint-modeling approach called *multidms*^48^. This open source package models two or more independent DMS experiments within the same predictive space, and utilizes appropriate forms of parameter regularization with the goal of identifying shifts in mutational effect We inferred shifts as the difference in abundance score in CYP2C19 relative to CYP2C9, such that positive shifts indicate a higher abundance score in CYP2C19 and vice versa. We tested regularization weights ranging from 0.0 to 1e-4, and selected 1e-5 as the optimal value for subsequent analysis (**Supp.** Fig. 6-8**, see Methods**). 2,366 variants (50.7%) had non-zero-shift values meaning that they had different effects between the two homologs, though most shifts had small effects (**Fig. 3B, Supp.** Fig. 9A).

**Figure 3.**
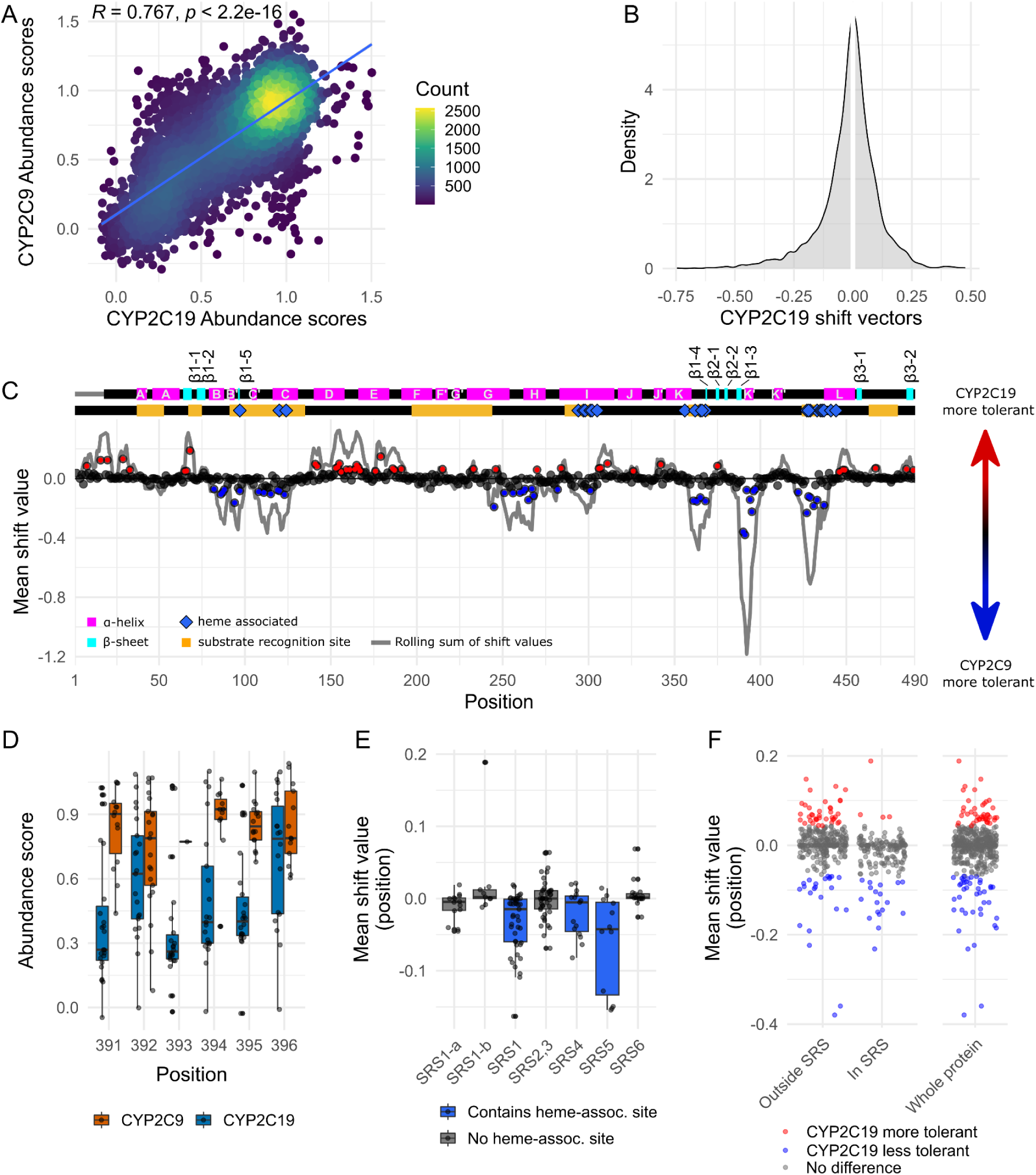
Comparison of VAMP-seq mutational tolerance of CYP2C19 and CYP2C9. A) Scatterplot of 5,979 abundance scores present in CYP2C19 and our previous CYP2C9 VAMP-seq experiment^30^. B) Distribution of non-zero shift values calculated using *multidms*^48^. Shift values are shown only for variants present in both datasets. C) Top: Secondary structure of CYP2C19 represented with α-helices shown in magenta and β-sheets shown in cyan. Middle: Substrate recognition regions are shown in orange, and sites that interact with heme are shown with blue diamonds. Bottom: Scatter plot of mean shift values for each position. Filled dots represent positions significantly more tolerant in CYP2C19 (red) or more tolerant in CYP2C9 (blue) with FDR-controlled p-values < 0.05 using a randomization test. The rolling sum of the mean shift values is depicted by the gray line. D) Boxplot of variant abundance scores for CYP2C19 (blue) and CYP2C9 (orange) across positions in the K’ helix. Dots represent variant abundance scores. E) Boxplot of shift values in substrate recognition sites (SRSs) with or without a heme-associated site. Dots represent mean shift values at each position within the SRS. F) Dot plot of mean shift values for each position separated by whether the position is in an SRS. Colors represent mean shift values that are significantly more tolerant in CYP2C19 (red), more tolerant in CYP2C9 (blue), or are not significantly different (gray) by randomization test.

We calculated the mean of the shift values at each position to reveal the effect of regional and structural features (**Fig. 3C**). We identified positions with mean shift values that differed significantly from 0 using a randomization test (**Supp.** Fig. 9B). The region with the largest mean shift values was in the K’ helix, which is part of a region that is both highly mobile and critical for packing of the hydrophobic core^5,49^ (**Fig. 3C**). In this region, mean shift values were negative, meaning that substitutions were more deleterious in CYP2C19 than in CYP2C9 (**Fig. 3D, Supp.** Fig. 10A).

Overall, CYP2C19 was more mutationally tolerant than CYP2C9 in the D, E, I, L, J, and J’ helices (**Fig. 3C, Supp.** Fig. 9A**, Supp.** Fig. 10A**, Bi-ii**), which form the majority of the hydrophobic core^5,49^. The sites that were most differentially tolerant in these helices were on portions of the helices that sit outside of the hydrophobic core (**Supp.** Fig. 10B**i-ii**). Conversely, CYP2C9 was more mutationally tolerant than CYP2C19 at positions within the hydrophobic core near important sites for heme positioning and function (**Supp.** Fig. 10A). Many of these heme-associated positions reside within substrate recognition sites (SRSs)^50^, and CYP2C9 was more mutationally tolerant than CYP2C19 in SRSs relative to the other regions of the protein (**Fig. 3C, E-F**). The mutational tolerances of positions in SRSs that were not heme-associated were similar between the homologs (**Fig. 3F**).

We also examined whether differences between CYP2C19 and CYP2C9 could be explained by sensitivity to variants of different biophysical types or by differences in the structures of the two homologs. However, we found that neither homolog is more sensitive to particular types of substitutions **(Supp.** Fig. 11A), and that shift values were unrelated to the distance between positions in the CYP2C19 and CYP2C9 crystal structures (**Supp.** Fig. 11B) Thus, comparison of variant effects between CYP2C19 and its closest homolog CYP2C9 revealed that CYP2C19’s K’ helix and, to a lesser extent, heme-associated positions in the hydrophobic core were more sensitive to mutation than CYP2C9, but that CYP2C19 was less sensitive than CYP2C9 to substitutions in other regions flanking the hydrophobic core.

### Amino acid swaps reveal homolog-specific constraints on abundance at sites influencing substrate specificity

We investigated abundance shifts at all variants comparing CYP2C19 and CYP2C9. However, the phenotypic differences in substrate recognition, membrane interaction, and thermodynamic stability between the two homologs must be driven by divergent sites. While most divergent sites are not localized to the catalytic site, some are critical for substrate specificity, regiospecificity, and stereospecificity^51–56^. In many cases, evolutionary pressures result in a protein’s reduction of thermodynamic stability in exchange for new functionality^31^. We wondered whether we could link the differences in thermodynamic stability between the homologs to substrate specificity by measuring the protein abundance of the 43 divergent sites. Thus, we investigated the abundance of the variants that partially convert CYP2C19 to CYP2C9 and vice versa.

We had abundance scores for 33 of the 43 2C19 variants that install the WT CYP2C9 amino acid, (e.g. CYP2C19→CYP2C9). For an exhaustive analysis, we individually measured GFP:mCherry fluorescence for each of the 10 CYP2C19→CYP2C9 variants not present in our abundance data (**Fig. 4A**). All but three CYP2C19→CYP2C9 substitutions were well tolerated. R261Q and L295F caused modest loss of abundance. Position 295 is critical for the specificity of CYP2C19 for S-mephenytoin and for the specificity of CYP2C9 for diclonefac^45,57^ whereas R261Q has not been studied in either homolog. V288E caused the largest loss of abundance of all of the swaps and was classified as “nonsense-like” (**Fig. 4A**). Position 288 alone does not have a known functional role. However, positions 241, 288, and 289 together have been suggested to play a role in abundance and substrate specificity^45,51,52,54,57^. In the CYP2C9 structure, K241 interacts electrostatically with E288 and hydrogen bonds with N289 to stabilize a region of the SRS4 in the I helix^51,56^. In CYP2C19, all three positions have different amino acids, E241, V288, and I289, and thus no electrostatic interaction between E241 and V288. Thus, the loss of abundance caused by V288E in CYP2C19 was likely due to the introduction of an electrostatic clash between E241 and V288E (**Fig. 4B**). Since all three positions contribute to functional differences between CYP2C19 and CYP2C9, we sought to understand how positions 241, 288, and 289 might interact to influence abundance.

**Figure 4.**
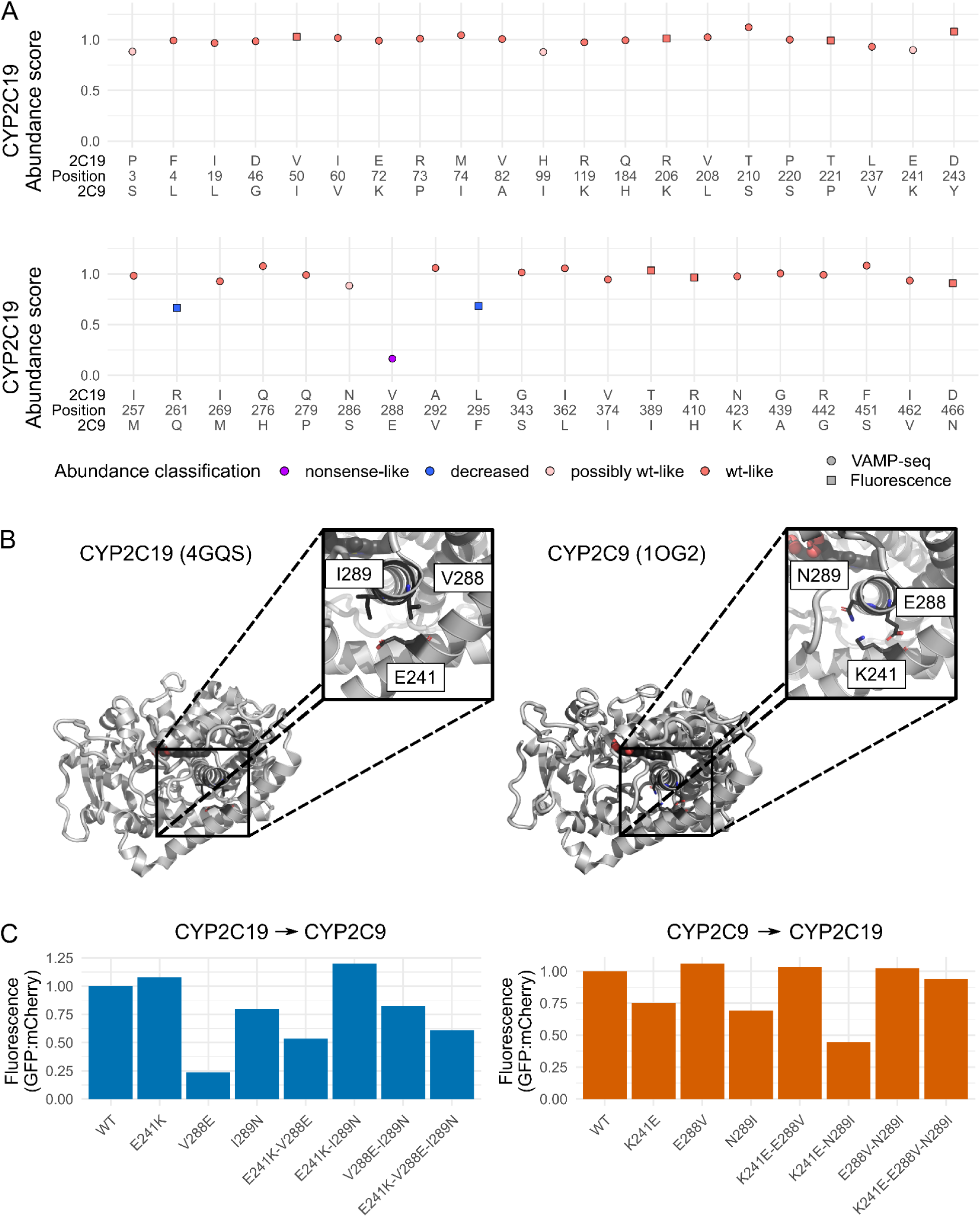
Abundance of CYP2C19 to CYP2C9 WT amino acid swaps. A) Dot plot of CYP2C19 abundance scores at positions that differ between CYP2C19 and 2C9. Each variant represents the abundance of CYP2C19 with the 2C9 WT amino acid installed. The WT amino acids for CYP2C19 and 2C9 are shown above and below the position. Dots are colored by 2C19 abundance classification as shown in the legend. Circles represent abundance scores derived from the VAMP-seq, and squares are individual GFP/mCherry fluorescence measurements normalized to CYP2C19 WT. B) CYP2C19 (PDB: <u>4GQS</u>) and CYP2C9 (PDB: <u>1OG2</u>) crystal structures. Positions 241 and 288 are shown as sticks and elements are colored (carbon: black, nitrogen: blue, oxygen: red). C) Bar plot of individually measured GFP/mCherry fluorescence for CYP2C19 (blue) and CYP2C9 (orange) variants. Each sample represents the geometric mean of GFP/mCherry fluorescence (n = 50,000 cells). Fluorescence of each variant is normalized to its respective WT CYP.

First, we individually measured the abundance of single and double mutants at positions 241 and 288 for both CYP2C19→CYP2C9 and CYP2C9→CYP2C19 variants (**Fig. 4B, C**). When individually measured, E241K had no effect on CYP2C19 abundance and V288E profoundly reduced abundance, the same effects we measured using VAMP-seq (**Fig. 4C**). Combining E241K and V288E partially restored CYP2C19 abundance. CYP2C9 K241E only modestly reduced abundance, even though this variant putatively results in an electrostatic clash similar to the one that dramatically reduced CYP2C19 abundance. CYP2C9 E288V had no effect on abundance, suggesting that the native K241-E288 electrostatic interaction likely does not contribute appreciably to thermodynamic stability^51^. Combining K241E and E288V fully restored CYP2C9 abundance (**Fig. 4C**). Thus, both homologs have a similar pattern, with installation of a second negative charge disrupting abundance. Elimination of one of the two negative charges even with variants from the other homolog restored abundance, although to differing degrees in each homolog.

Next, to incorporate position 289 into our analysis, we measured the abundance of the CYP2C19→CYP2C9 and CYP2C9→CYP2C19 241, 288, 289 triple mutants (**Fig. 4C**). The CYP2C19→CYP2C9 triple mutant had a modestly reduced abundance relative to CYP2C19 WT, largely restoring the low abundance of the E241K, V288E double mutant. The CYP2C9→CYP2C19 triple mutant had an abundance equivalent to CYP2C9 WT and to each of the two double mutants. Thus, while the interaction between these three positions is complex, it seems likely that sequence changes in this region of the protein contribute to the increased thermodynamic stability of CYP2C19.

### Annotating human *CYP2C19* variants

CYP2C19 variants can increase, decrease, or eliminate an individual’s ability to metabolize many important drugs, and knowing variant function can help avoid severe and expensive adverse events^13–15,58,59^. For example, the anti-platelet drug clopidogrel is activated by CYP2C19. Thus, individuals with deleterious CYP2C19 variants experience reduced or non-existent benefit from clopidogrel, requiring higher doses or alternative drugs. Genetic testing for CYP2C19 variants prior to clopidogrel treatment is important for avoiding major adverse cardiovascular events^19,20^. PharmVar is a repository for pharmacogene allelic variation and functional information, including *CYP2C19*. Alleles in PharmVar are known as “star alleles,” and annotated using star notation^17^. For example *CYP2C19*5* refers to R433W. Despite decades of study, 10 of the 39 *CYP2C19* star alleles are of uncertain function. Thus, we analyzed the functional effects of *CYP2C19* alleles in PharmVar. All four PharmVar “normal function” alleles had WT-like abundance scores (**Fig. 5A**). Of the eight “decreased” and “no function” alleles, six were low abundance. The remaining two decreased/no function alleles, *6 and *9, were WT-like abundance. PharmVar lists the *6 allele (R132Q) as “no function” with “definitive” evidence; however we measured a WT-like abundance score of 1.06 (95% CI 1.09 – 1.03). This strongly suggests that the *6 allele’s loss of function results from disrupted enzymatic activity rather than loss of abundance. Consistent with this interpretation, the *6 allele is intact enough to bind its heme cofactor, but has a decreased ability to metabolize substrates and disrupted electron flow from cytochrome P450 reductase^60,61^. Allele *\**9 (R144H) had an abundance score of 0.914 (95% CI 0.956-0.872). Consistent with these results, *9 has WT-like affinity for cytochrome P450 reductase^62^, suggesting that it is at least partially folded. However, our abundance results conflict with another, smaller scale VAMP-seq experiment in which *CYP2C19*9* was identified as “decreased” abundance^34^ with less than ∼50% of WT abundance. We therefore individually validated this variant and reaffirmed its WT-like abundance in our hands (**Supp.** Fig. 12). In light of *9’s moderately reduced activity against mephenytoin, ability to bind cytochrome P450 reductase normally and WT-like abundance in our assay, we suggest that, like *6, *9 has normal abundance but decreased catalytic activity. Overall, six of the eight known loss of function alleles had reduced abundance, and all normal function alleles had WT-like abundance. While variants with WT-like abundance could have low or no function, reflecting the many ways function can be compromised, low abundance variants were always low or no function alleles. Thus, abundance is a powerful method for identifying loss of function variants. Of the ten PharmVar alleles with uncertain function, four were present in our VAMP-seq library, and we found that *30 and *23 had decreased abundance strongly suggesting that they would disrupt drug metabolism (**Fig. 5A**).

**Figure 5.**
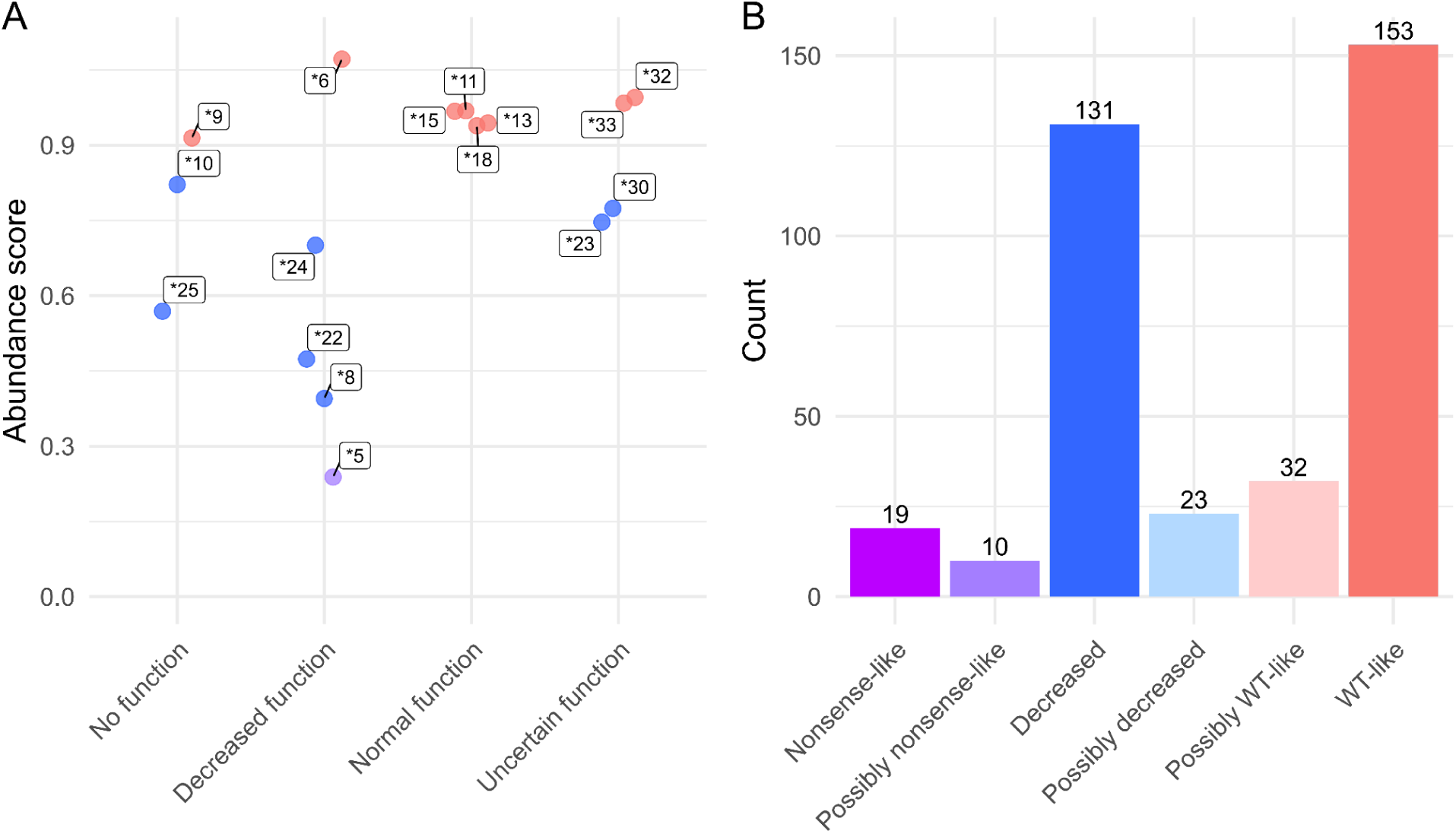
CYP2C19 abundance scores for variants found in humans. A) Scatter plot of CYP2C19 abundance scores of star (*) alleles with clinical functional status according to CPIC database (accessed May 18, 2022). Dots are colored by abundance score classification and labeled by their star allele designation. B) Bar plot representing abundance score classification of variants in gnomAD (accessed May 18, 2022).

As sequencing and genetic testing are more widely deployed, rare variants with unknown clinical consequences are being identified at an exponentially increasing rate^63^. Reflecting this reality, the annotated alleles in the PharmVar database are only a fraction of all of the CYP2C19 variants discovered so far. There are 408 unique *CYP2C19* missense variants in the exome database gnomAD, 390 of which have no CPIC annotation or functional information. We annotated 368 (90.2%) of the variants in gnomAD (**Fig 5B**) and identified 131 (35.6%) variants with “decreased” abundance and 29 (7.88%) with “nonsense-like” or “possibly nonsense-like” abundance relative to WT, strongly suggesting that these variants have decreased or no function. We annotated 210 (57.0%) variants as “WT-like” or “possibly WT-like,” indicating that these variants may have normal function. However, assessment of enzymatic activity would be needed to definitively determine if these “WT-like” or “possibly WT-like” variants have normal function since variants can eliminate activity without affecting protein stability. These results are broadly consistent with a study that genotyped 2.29 million participants for *CYP2C19*\*2, *3, and *17 alleles. The study discovered that *2 was present in 15.2%, *3 in 0.3%, and *17 in 20.4% of individuals, and nearly 60% had at least one of these star alleles^64^. Thus, *CYP2C19* variants with reduced abundance appear common in the population.

## Discussion

The CYP family tree spans all animal kingdoms and comprises an exceptionally versatile set of enzymes. Understanding the phenotypic consequences of natural variation in human CYPs is particularly important since they catalyze the metabolism of most drugs currently in use. However, even closely related CYPs, like CYP2C19 and CYP2C9, are functionally distinct, and the underlying causes of these distinctions are largely unknown. We used VAMP-seq to measure the abundance of 7,660 CYP2C19 missense variants. In addition to confirming positions known to be critical for CYPs structure and function, we revealed that variants at four conserved positions in the hydrophobic core do not impact CYP2C19 abundance. By jointly analyzing 4,670 shared CYP2C19 and CYP2C9 abundance scores, we discovered regions where the two homologs have different mutational tolerances. CYP2C9 has a more tolerant hydrophobic core, whereas CYP2C19 is more tolerant in regions surrounding the core. We measured the abundance of WT amino acid swaps between CYP2C19 and CYP2C9, discovering a region likely responsible for at least some of the thermodynamic stability difference between the homologs. Finally, our abundance scores identify known reduced activity CYP2C19 variants with high fidelity, and indicates that two star alleles of unknown function, *30 and *23, likely have reduced abundance. We also evaluated 368 of the 408 human CYP2C19 variants with no prior annotation. Notably, 43% of these variants are low abundance, warranting follow-up studies to measure their impacts drug metabolism.

Of the 58 positions conserved across eukaryotic CYPs, 52 had more than 65% reduced abundance variants when substituted with amino acids of a different biophysical type. The remaining six were surprisingly mutationally tolerant. The conservation and tolerance of positions 136 and 322 can be explained by their location on the surface of the protein, and are likely involved in binding cofactor cytochrome P450 reductase (CPR), as they do in the closely related CYP2C9^39,40^. However, positions, 297, 300, 301, and 362, were tolerant to mutations despite being in the hydrophobic core where mutations are nearly always deleterious. These positions impact substrate specificity of many drugs in CYP2C9 including warfarin, flurbiprofen, and acetaminophen^7,65,66^. The significance of these positions in CYP2C19 lies not only in their impact on substrate specificity but also in their specialized role, as they primarily influence specific functions rather than overall protein abundance.

We also jointly analyzed CYP2C19 and CYP2C9^30^ abundance scans using *multidms*^48^ to find variants with different abundances. Variants in the K’-helix reduced abundance in CYP2C19 but were tolerated in CYP2C9, suggesting a markedly different mutational tolerance of K’ in the enzymes despite having identical WT amino acids. Moreover, CYP2C9 had a more tolerant hydrophobic core than CYP2C19, especially in SRSs that contain heme-associated positions. CYP2C9’s higher mutational tolerance in its core may indicate more flexibility. We speculate that, since the flexibility of CYP active sites is correlated with its promiscuity^44,67^, this may allow CYP2C9 to bind more substrates^9^.

Variants capable of imparting novel function, like those that alter substrate specificity, often reduce thermodynamic stability^31^. To determine if CYP2C9’s altered substrate profile and lower thermodynamic stability relative to CYP2C19^11^ constituted such a tradeoff, we analyzed the 43 divergent positions between CYP2C19 and CYP2C9. We found that positions 241, 288, and 289 are the likely locus of such a tradeoff because these three positions impact substrate specificity^51,52,54^ and they are also adjacent in the structure of both enzymes. Position 288 was the only position in CYP2C19 where installing the CYP2C9 amino acid caused profound loss of abundance, and combining it with swaps at 241 and 289 revealed that these sites have distinct interactions in each homolog. Thus, we hypothesize that these positions are partially responsible for the difference in thermodynamic stability. The CYP2C19 V288E substitution likely causes loss of abundance because it places a negative charge adjacent to E241. Likewise, CYP2C9 K241E introduces the same opposing negative charge adjacent to E288 and also reduces abundance. This pattern of loss of abundance suggests that CYP2C9 may have evolved from CYP2C19. This is because CYP2C9 is the only enzyme in the family that has a negatively charged amino acid at 288, meaning that the ancestral sequence had valine at position 288^56^. Thus, the ancestral CYP2C9 likely acquired E241K or I289N first, both of which partially ameliorate the loss of abundance induced by V288E. We did not find that any combination of swaps at these three positions could fully restore CYP2C19 V288E abundance. One possibility is that swaps at other sites, which by themselves do not affect CYP2C19 abundance, could fully rescue V288E. However, the reduced abundance of the CYP2C19 E241K-V288E-I289N variant is in line with CYP2C9’s reduced thermodynamic stability. CYP2C9’s new substrate binding capabilities apparently made this loss of thermodynamic stability evolutionarily tolerable.

Importantly, our VAMP-seq derived abundance data has limitations. First, the abundance of each variant arises from the balance between protein synthesis and degradation driven by cellular protein quality control systems. In VAMP-seq, all variants are expressed transgenically from an inducible promoter, and an mCherry reporter expressed via an IRES is used to control from cell-to-cell variation in expression. Thus, VAMP-seq derived abundance scores largely reflect changes in degradation which are, in turn, generally driven by stability-related changes in protein folding^22,24,28,29^. However, abundance changes can result from other mechanisms such as changes to degron sequences or protein localization. Second, we expressed the CYP2C19 cDNA from an inducible promoter, meaning we cannot detect variants that induce splicing defects or affect transcriptional regulation. Third, variants can affect function without affecting abundance. For example, variants may disrupt a critical substrate binding position or prohibit binding to critical cofactors like cytochrome P450 reductase (CPR) or cytochrome b5. Therefore, while variants that we identified with reduced abundance are likely to alter drug metabolism, variants with WT-like abundance may not necessarily have normal function. Finally, the VAMP-seq assay depends on fluorescent reporters and fluorescence activated cell sorting. As a result, subtle changes in abundance are difficult to discern.

In the future, we envision intersecting mutational scans from important CYPs in other subfamilies such as CYP2D6 and CYP3A4. Since *multidms* is capable of jointly analyzing more than two scans, it will be a powerful tool to compare mutational effects across additional CYPs, helping us understand the extent to which variant effects are conserved across the family. In addition to improving personalized drug dosing, such comprehensive profiling could expand our overall understanding of CYP function.

## Methods

### General reagents

Unless otherwise noted, all chemicals were obtained from (MilliporeSigma) and all the enzymes were obtained from New England Biolabs. All cell culture reagents were purchased from Thermo Fisher unless otherwise noted. All plasmids and oligonucleotides used in this study are listed in Table S3.

### Growth media and culturing techniques

HEK293T cells (ATCC CRL-3216) and the derived landing pad cell line were cultured in Dulbecco’s modified Eagle medium supplemented with 10% fetal bovine serum, 100U/mL penicillin, and 0.1 mg/mL streptomycin. Landing pad expression was induced with doxycycline at a final media concentration of 2.5 μg/mL. Cells were passaged by by detachment with trypsin 0.5%. All cell lines were tested negative for mycoplasma on a monthly basis.

### Library mutagenesis

The CYP2C19 library was constructed using inverse PCR-based site-directed saturation mutagenesis^68^. Saturation mutagenesis primers were designed for each codon of *CYP2C19* across positions 2 through 490. Each forward primer contained an NNK at the 5’ end of the sequence. Primers were obtained from Integrated DNA Technologies (IDT). Our library consisted of 7,660 of 9,291 (82.3%) possible missense substitutions represented by 147,723 unique barcodes (mean of 11.87 and median of 7 for single amino acid variants; see Table S2 for details).

CYP2C19 WT was codon optimized for human expression in a pHSG298 backbone. We completed inverse PCRs using NNK oligos for each position excluding the methionine at position 1. Each PCR reaction contained 125 pg of template, 2uM of mixed primers, and 5% DMSO in a 5 μl reaction volume of KAPA HiFi Hotstart 2x ReadyMix. The resulting products were confirmed by visualizing on a gel and quantified using either the Quant-iT PicoGreen dsDNA Assay Kit (Invitrogen) or Qubit fluorometry (Life Technologies). The PCR products were then pooled at equimolar ratios and cleaned using the DNA Clean and Concentrator Kit (Zymo Research), followed by gel extraction. The pooled libraries were 5’ phosphorylated with T4 polynucleotide kinase and subjected to intramolecular ligation overnight. Next, 8.5 μl of phosphorylated product was combined with 1 μl of 10x T4 ligase buffer and 0.5 μl of T4 DNA ligase (NEB), incubated at 16℃ overnight, and cleaned and concentrated. The ligated products were transformed into electrocompetent E. coli cells (NEB C2989K or C3020K) with electroporation at 2 kV, and the resulting transformants were plated on LB + kanamycin. The CFUs on the plates were counted to estimate the unique molecules transformed and to estimate the coverage of the library. Finally, the library was subcloned into the expression and recombination vectors and barcoded.

To generate barcoded libraries, the variant library was first digested with SacII and AflII at 37℃ for 1h, followed by heat inactivation at 65℃ for 20 minutes. We ordered barcode oligos with 18bp random sequences from IDT, resuspended them at 100 μM, and annealed them by combining 1 μl of each primer with 4 μl of CutSmart buffer and 34ul of ddH2O and running 98℃ for 3 minutes, ramping down to 25℃ at –0.1 ℃/s. The annealed oligos were then Klenow filled by combining 0.8 μl Klenow polymerase (exonuclease negative, NEB) with 1.35 μl of 1mM dNTPs with 40 μl of product to fill in the barcode oligo, using cycling conditions of 25℃ for 15 min, 70℃ for 20 min, ramping down to 37℃ at –0.1℃/s. The resulting products were then ligated overnight at 16℃. The barcoded library was transformed into electrocompetent E. coli cells (NEB C2989K) and midiprepped (QIAGEN). The size of the barcoded library was bottlenecked and estimated by colony counts to be 67,000.

To obtain more accurate library counts, we sequenced the libraries with Illumina sequencing. The forward and reverse reads were merged using Pear^69^, and barcode counts were estimated using Bartender^70^. Barcodes with fewer than 10 reads were filtered out, resulting in ∼200,000 unique barcodes for an average of 21x coverage.

### PacBio sequencing for barcode-variant mapping

PacBio sequencing libraries were generated with SMRTbell Express Template Prep Kit 3.0 (Pacific Biosciences) according to manufacturer’s instructions. The barcoded variant sequences were excised using restriction enzymes NheI-HF and HindIII-HF and purified with AMPure PB beads (Pacific Biosciences 100-265-900) at a 1:1 ratio of beads to DNA. Following end-repair, A-tail attachment, and ligation, the assembled product was extracted using a BluePippin instrument (Sage Science, BLU0001) using a 0.75% agarose precast cassette (Sage Science, BLF7510). Library purity and size was confirmed by 4200 TapeStation (Agilent, G2991BA) before sequencing. Samples were submitted to University of Washington PacBio Sequencing Services and sequenced on one SMRT cell in a Sequel II v2.0 run using a 15 hour movie.

We filtered long reads for a minimum of 3 passes. We then analyzed the circular consensus reads (CCSs) using PacRAT to identify and link the gene variants with the barcode region^71^. The filtered barcode-variant library contained 12,559 unique nucleotide sequences tagged by 176,372 unique barcodes (see Supplemental Table S3 for details).

### FACS-based deep mutational scan (VAMP-seq)

HEK293T with a Bxb1 serine recombinase landing pad with an inducible Caspase 9 cassette (HEK293T-LLP-iCasp9)^33^ that enable expression of one variant per cell were used for all human cell experiments. To recombine the variant library into HEK293T cells, 3,500,000 cells were seeded in 10 cm plates (2-4 per replicate) and transfected with FuGENE® 6 Transfection Reagent (Promega, E2692). In one tube, 7.1 μg of barcoded library plasmid was mixed with 0.48 μg of Bxb1 plasmid in 710 μL of OptiMEM. In a separate tube, 28.5 μl of Fugene was diluted into 685 μl of OptiMEM. The Fugene and DNA tubes were then combined and incubated at room temperature for 15 minutes. The Fugene/DNA mixture was added to cells dropwise, and cells were incubated for a minimum of 48 h before induction with doxycycline at a final concentration of 2.5 μg/mL. 24 h after doxycycline was added, we added AP1903 at a final concentration of 2 nM to induce Caspase 9 dimerization and eliminate all unrecombined cells.

Transfected HEK293T cells were sorted using a BD AriaIII sorter. Cells were gated for live, recombined singlets. In recombined cells, the ratio of GFP:mCherry fluorescence was calculated and plotted as a histogram. The histogram was split into four quartiles. Each quartile was sorted into separate 5 mL tubes. Cells from each bin were grown out for 1-2 days to ensure enough DNA for sequencing. Three biological replicates from separate transfections were collected for the FACS-based deep mutational scan.

### Sorted abundance library amplification and sequencing

Sorted cells were harvested and pelleted by centrifugation, and then stored at –20°C until all replicates were collected. Genomic DNA was extracted using the DNEasy Kit (QIAGEN) according to the manufacturer’s instructions, with the addition of a 30-minute incubation step at 37°C with RNase during the resuspension step. For the first round of PCR, eight 50 μL reactions were set up for each sample, with a final concentration of 50 ng/μL input genomic DNA, 1x Q5 High-Fidelity Master Mix, and 0.25 μM of JS454 and JS1004 primers. The reaction conditions were 95°C for 30 seconds, 98°C for 10 seconds, 60°C for 30 seconds, 72°C for 3 minutes, repeated 4 additional times, followed by 72°C for 2 minutes and a 4°C hold. The eight reactions were then combined, bound to AMPure XP (Beckman Coulter) at 0.6X bead volume to sample volume, cleaned, and eluted with 38.5 μL water. 15 μL (40%) of the eluted volume was mixed with Q5 High-Fidelity Master Mix, GB001, and one of the indexed reverse primers, JS385 through JS473, added at 0.25 μM each. The PCR reaction was run with SYBR Green I on a Bio-Rad MiniOpticon. The reaction was denatured for 3 minutes at 95°C, cycled 18 times at 95°C for 15 seconds, 67°C for 30 seconds, and 72°C for 45 seconds, with a final 2-minute extension at 72°C.

The indexed amplicons were then run on a Tapestation according to the manufacturer’s instructions. For each sample, 1 μL sample was mixed with 3 μL of Sample Buffer, thoroughly mixed, and run on a D1000 ScreenTape (Agilent Technologies) using an internal electronic ladder. The bands were quantified using the TapeStation analysis software. The samples were then pooled in equal amounts, loaded onto a 1% agarose gel with SYBR Safe, and then the gel was extracted using a freeze and squeeze column (Bio-Rad). Finally, the quantification of the pooled sample was done with the Qubit™ 1X dsDNA Assay Kit broad range (Q33266).

### Library sequence analysis

Barcode sequences were trimmed and filtered for a minimum base quality of Q20 using the FASTX-toolkit. These barcodes were then used to generate a FASTQ file input for Enrich2 to count variants. Variants with insertions, deletions, or multiple amino acid substitutions were excluded. Barcode counts were then collapsed to variant counts, retaining variants with a total frequency greater than 4 x 10^-5 across all bins (**Supp.** Fig. 2). For each replicate, an abundance score was calculated using a weighted average of variant frequency across bins (*w*_1_ = 0.25, *w*_2_ = 0.5, *w*_3_ = 0.75, *w*_4_ = 1)^22^. Scores were normalized to synonymous and nonsense distributions, excluding the top 20% of nonsense scores. Missense variant abundance scores ranged from –0.09 to 1.5.

Abundance classes were determined as in previous studies^22,30^. To discern between “WT-like” and “decreased” scores, we used a synonymous score threshold. This threshold was set at the 5th percentile of synonymous scores (0.856). Variants were classified as “WT-like” if their lower confidence interval exceeded the threshold, or as “possibly WT-like” if only their score surpassed the threshold. Additionally, an upper threshold at the 95th percentile of synonymous scores (1.14) was used to differentiate between “WT-like” and “increased” scores. To distinguish between “decreased” and “nonsense-like” scores, we used a threshold at the 95th percentile of nonsense scores (0.265). Variants were categorized as “nonsense-like” if both their score and upper confidence interval were below the nonsense threshold, or as “possible nonsense-like” if only their score fell below the threshold. Out of a total of 8,480 variants, 316 were nonsense, 504 were synonymous, and 7,660 were missense variants. The missense variants were categorized into the following abundance classes: 2,590 WT-like, 612 possibly WT-like, 3,146 decreased, 437 possibly decreased, 340 possibly nonsense-like, and 708 nonsense-like.

### VAMP-seq internal validation with individual variants

We cloned 11 variants using IVA cloning^72^ site directed mutagenesis into the VAMP-seq recombination vector (attB-CYP2C19-eGFP-IRES-mCherry) via primers listed in **Table S3** (HB049 through HB073, GB143, and GB144). Mutations were generated with KAPA HiFi DNA Polymerase (KAPA Biosystems KK2601) and 40 ng of *CYP2C19* template plasmid attB-CYP2C19-eGFP-IRES-mCherry. After completing inverse PCR for each variant, we digested the products with DpnI to eliminate remaining WT template, and transformed chemically competent *E. coli* cells (NEB C2987 or Bioline BIO-85027). Bacterial clones were prepped with a midiprep kit, validated by Sanger sequencing and whole plasmid nanopore sequencing. We then transfected the preps into and HEK293T-LLP-iCasp9 landing pad cells in 6-well plates with 400,000 cells per well. 2.7 μg of plasmid was mixed with 0.300 μg of Bxb1 plasmid in 125 μL of OptiMEM and 5 μL P3000 reagent. In a separate tube, 2.25 μL of Lipofectamine was added to 125 μL of OptiMEM. The tubes were then combined and incubated at room temperature for 15 minutes. After incubation, the Lipofectamine/DNA mixture was added to cells dropwise and the plates were placed in an incubator at 37℃. After 24 hours, the cells were induced with doxycycline at a final concentration of 2.5 μg/mL, and at least 24 hours later we selected for recombinant cells by adding small molecule AP1903 which causes inducible Caspase 9 in unrecombined landing pad cells to dimerize, activate, and induce apoptosis.

Recombined cells were grown to full confluence and analyzed with a BD LSRII flow cytometer. Cells were gated for live, recombined singlets. We calculated a ratio of eGFP/mCherry fluorescence, and the geometric mean of the distribution of this ratio was reported. Flow cytometry data were collected with FACSDiva V8.0.1 (BD Biosciences) and analyzed with FlowJo V.10.8.1 (Ashland, OR). Three biological replicates of each individual variant were measured.

### *multidms* analysis of CYP2C19 and 2C9 deep mutational scans

To analyze and compare the deep mutational scanning (DMS) data between the CYP2C19 and CYP2C9^30^ homologs, employed a novel approach to jointly model the mutational effects using an open source package, *multidms*^48^. Using highly multiplexed libraries we densely sampled many of the possible mutations across the two homologs. Variants within each library were limited to at most one mutation, yet each individual variant may have been associated with many unique barcodes (**Supp.** Fig. 13). To use *multidms*, we calculated abundance scores at the level of individual barcodes such that the model had to infer expected mutation effects of the CYP2C9 variants (β), while jointly identifying if (and by how much) the respective mutation effects may have shifted in the CYP2C19 background (*Δ*). The input data consisted of barcode-level abundance scores from two biological replicate DMS experiments for both homologs. We then grouped these four raw datasets into two independent training datasets, each of which consisted of one replicate from each CYP2C19 and CYP2C9. We then separately fit *multidms* models to the training sets and found the resulting fits to have well-correlated parameter sets (**Supp.** Fig. 6-7).

The models were trained to predict abundance scores in CYP2C9 and shifts in scores in CYP2C19 relative to CYP2C9, minimizing the difference in predicted and experimentally measured scores, as quantified by a Huber loss function. A key feature of the *multidms* modeling approach is the application of a lasso (L1) penalty to the shift parameters during the fitting procedure, effectively driving them to zero unless they are strongly supported by the data. To determine a reasonable penalty coefficient (λ) for the lasso, we compared the results of seven model fits – each using different coefficients ranging from λ=0, to λ=5e-4. This λ sweep was performed in duplicate using each of the distinct training sets, for a total of 14 model fits evaluated. **Supp.** Fig. 8 shows summary statistics of the model fits. This figure emphasizes the accuracy-simplicity tradeoff for any given value of λ. For example, as λ increases, we observe higher overall training set loss, but also increased correlation between replicate parameters values. We selected a penalty coefficient of λ = 1e-5 as a reasonable value to balance model accuracy with shift sparsity. The shifts in the main text are then reported as the averaged values between the replicate model fits, each using the selected penalty coefficient. See https://github.com/matsengrp/CYP-multidms for the code used for this analysis.

We identified positions whose mean shift value was significantly different from the distribution of all shift values using a randomization test. To generate a null distribution, we randomly sampled 10 shift values, the average number of abundance scores per position, and calculated the mean of the shifts. This procedure was repeated 100,000 times. We calculated p-values for each position by counting the number of randomly generated mean shifts more extreme than the position mean and dividing it by 100,000, the total number of randomly generated shifts. P-values were then adjusted for repeated hypothesis testing using the Benjamini-Hochberg method^73^ with a false discovery rate of 5%. Positions with p-values less than 0.05 were considered significant.

## Additional files

Table S1: CYP2C19 library fluorescence activated cell sorting

Table S2: Library statistics from barcode-variant mapping

Table S3: Plasmids and oligos used in this study

Table S4: CYP2C19 variant abundance scores

Table S5: CYP2C19 star allele CPIC functional annotations

Table S6: CYP2C19 individual variant validation of abundance scores

Table S7: *multidms* analysis of CYP2C9 and CYP2C19 abundance scores

## Data and code availability

The accession number for the sequencing data reported in this paper is NCBI GEO: GSE244489. Code and processed variant scores generated during this study are available at GitHub: https://github.com/FowlerLab/cyp2c19_2c9. *multidms* analyses and data are available at GitHub: https://github.com/matsengrp/CYP-multidms

**Supplemental Figure 1.**
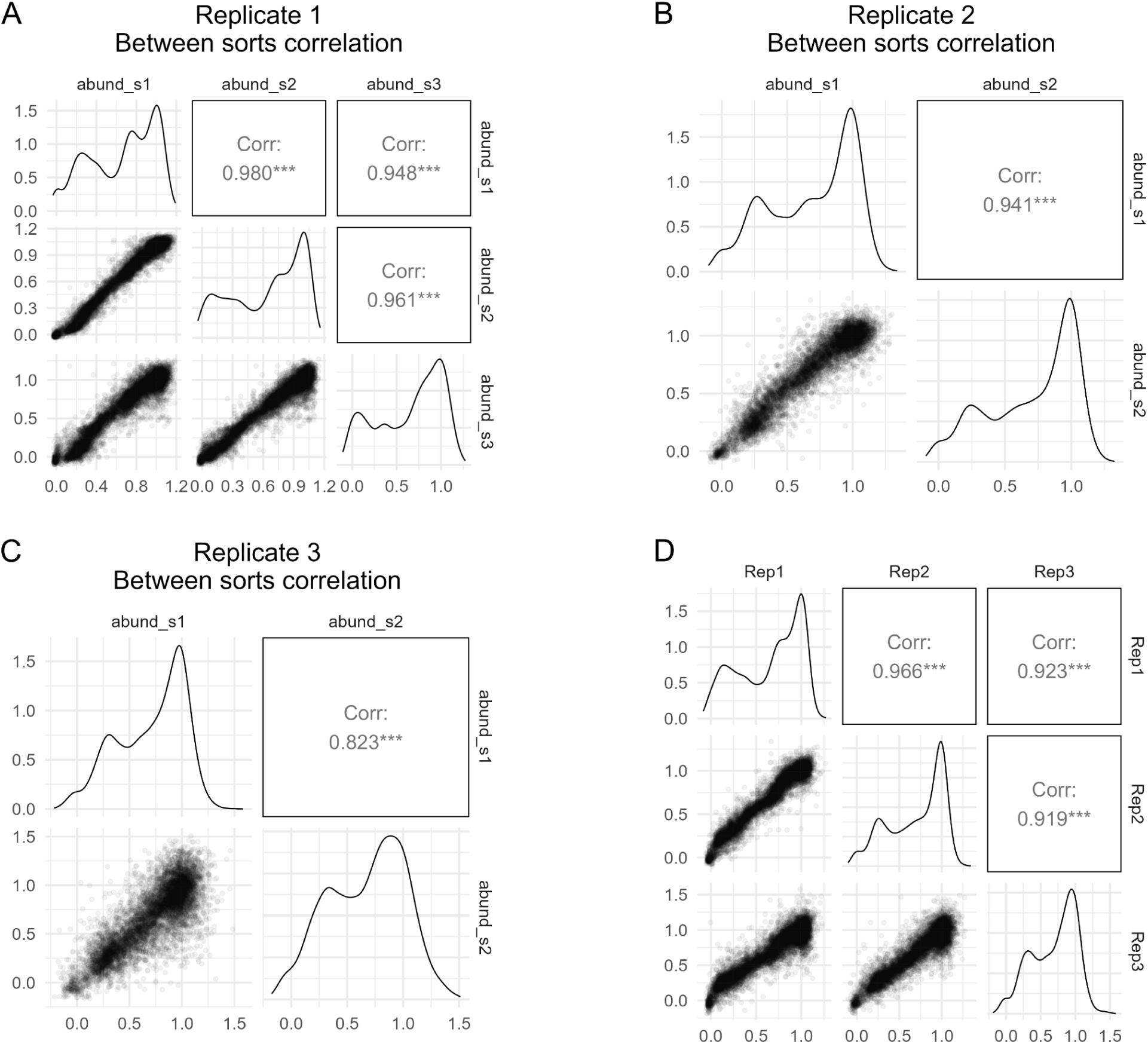
Abundance score correlation matrices. A-C) Scatter plots showing correlation between variant abundance scores between each sort for three replicates. D) Scatter plot showing correlation of variant abundance score across all replicates. Sorts were combined by averaging variant abundance scores.

**Supplemental Figure 2.**
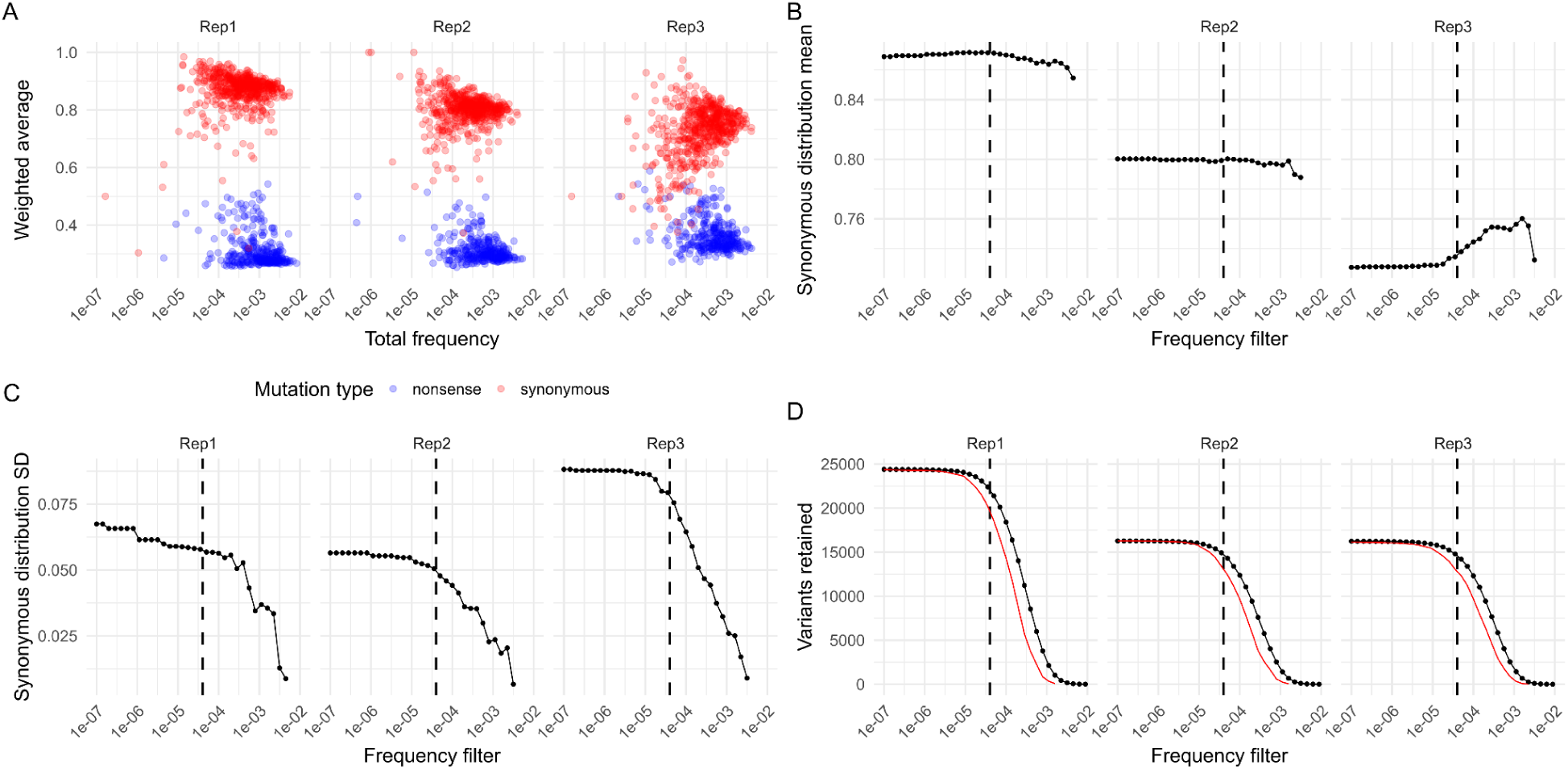
Determining variant frequency filters. A) Scatter plots of variant weighted averages and frequencies for synonymous (red) and nonsense (blue) variants across three replicates. B-D) Frequency filters for abundance scores. Dot plots show the synonymous distribution means (B), standard deviations (C), and number of variants (D; black) or 14 times the number of synonymous variants (red) for each replicate across frequency cutoffs. For plots E-H, the frequency filter of 4×10^-5^ used is shown as a dotted line.

**Supplemental Figure 3.**
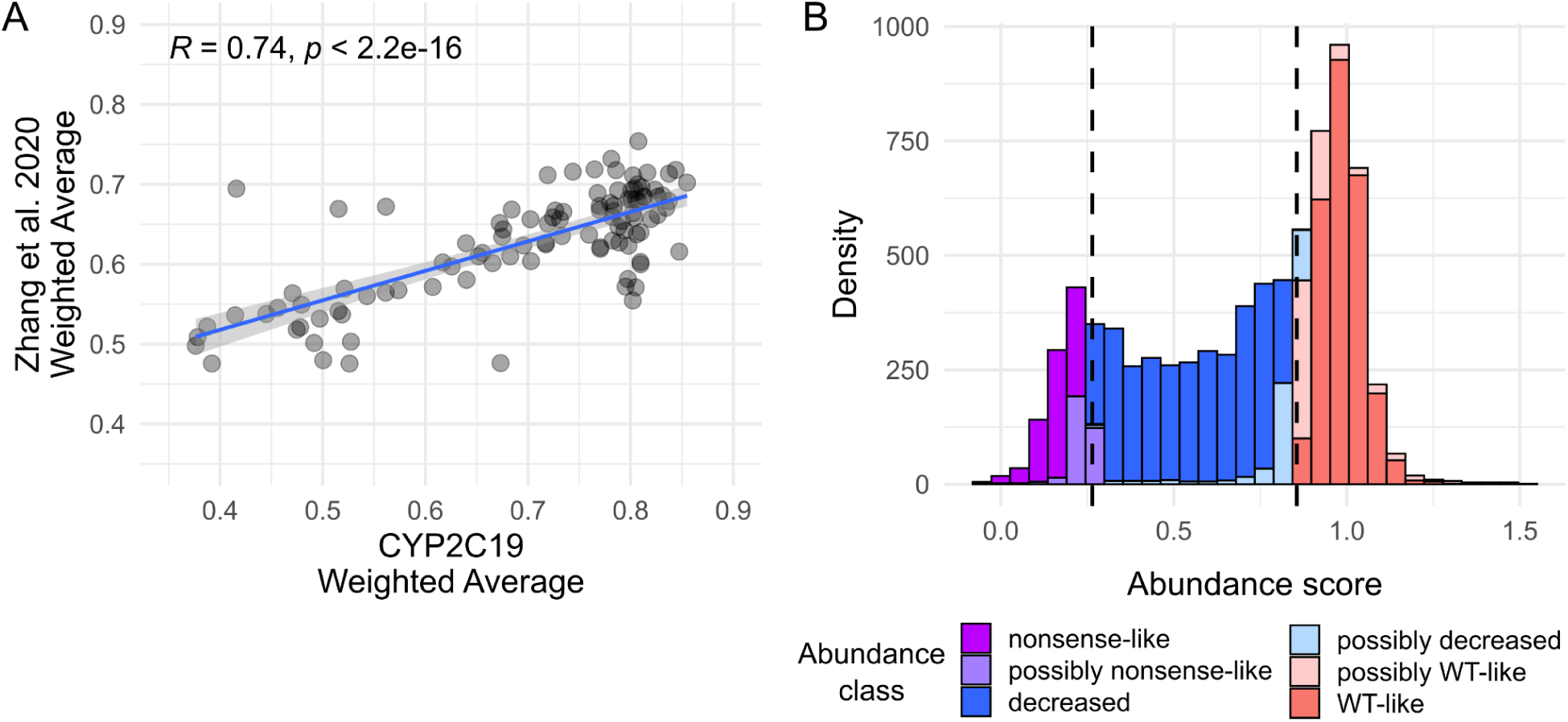
Categorization of variant abundance scores into classes. A) Scatter plot of CYP2C19 abundance weighted averages compared between our assay and another VAMP-seq assay completed by Zhang et al, 2020^34^. Pearson’s R = 0.74 B) Missense variant abundance score histogram colored by abundance class. In dotted lines, scores lower than the 5th percentile of the synonymous distribution (right) and 95th percentile of the nonsense distribution (left) were used for classification. Variant abundance class was determined by whether their scores and confidence intervals fell in the synonymous and nonsense distribution thresholds, as described in the Methods.

**Supplemental Figure 4.**
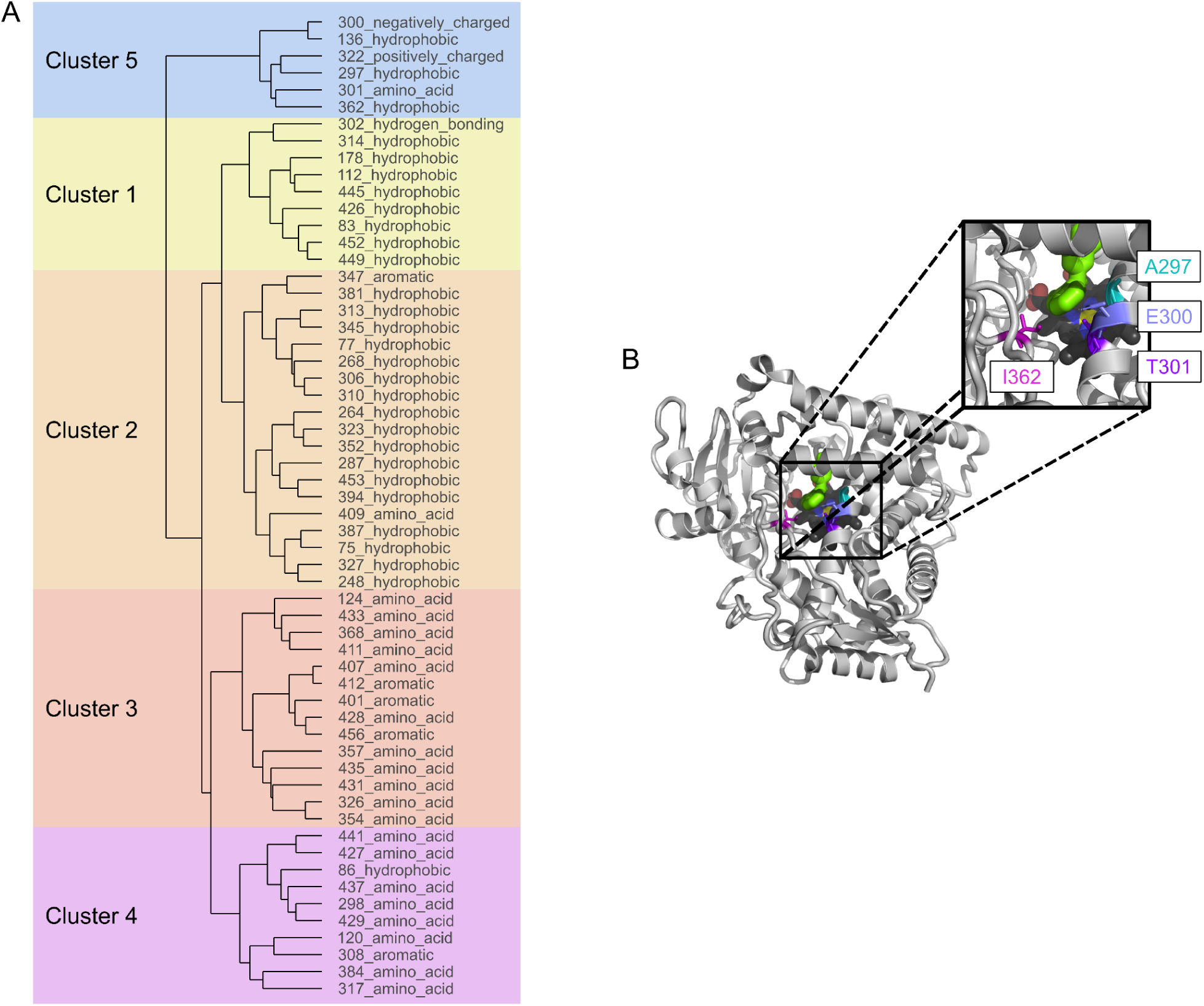
Mutationally tolerant conserved sites in the hydrophobic core. A) Dendrogram showing hierarchical clustering of sites conserved in >80% eukaryotic CYPs ^38^. B) CYP2C19 structure (PDB: <u>4GQS</u>). Heme is colored by element (carbon: black, nitrogen: blue, oxygen: red, iron: yellow), and PDB chemical 0XV in green. Positions A297 (cyan), E300 (lavender), T301 (purple), and I362 (magenta) are located in the hydrophobic core near heme.

**Supplemental Figure 5.**
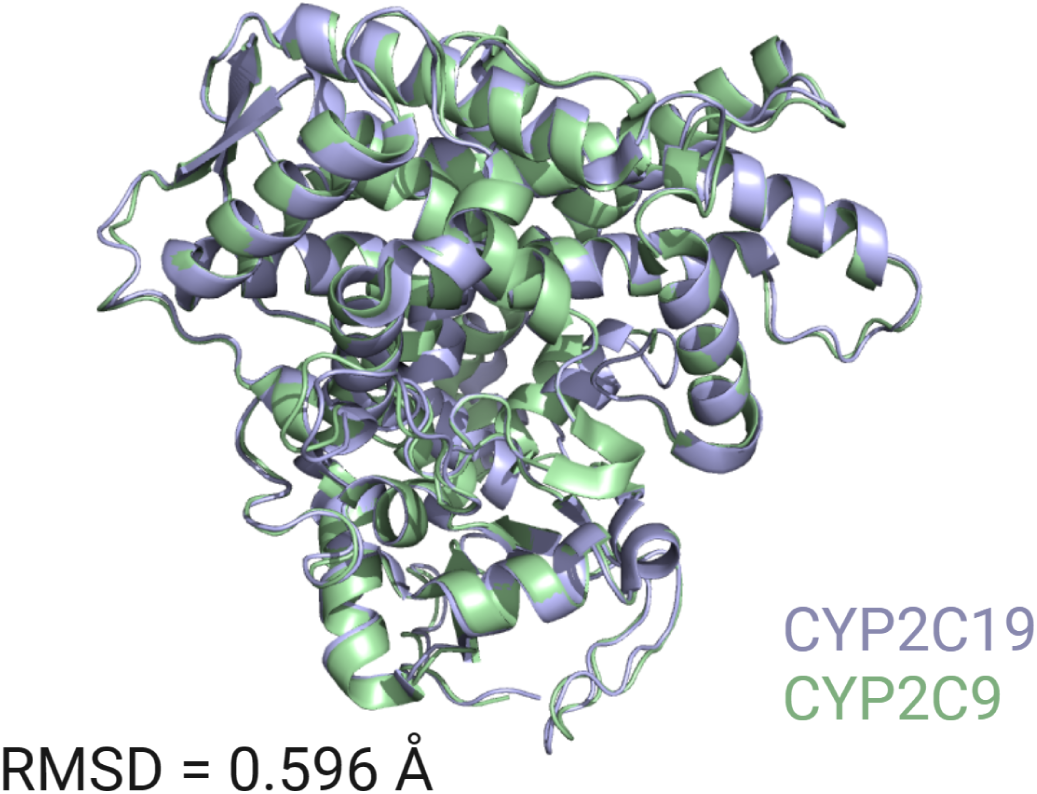
CYP2C19 and CYP2C9 crystal structure similarity. Overlaid cartoon representations of CYP2C19 (PDB: <u>4GQS</u>; purple) and CYP2C9 (PDB: <u>1OG2</u>; green) crystal structures. Alignment generated using PyMOL. Root mean-squared deviation of backbone (RMSD) = 0.596 Å.

**Supplemental Figure 6.**
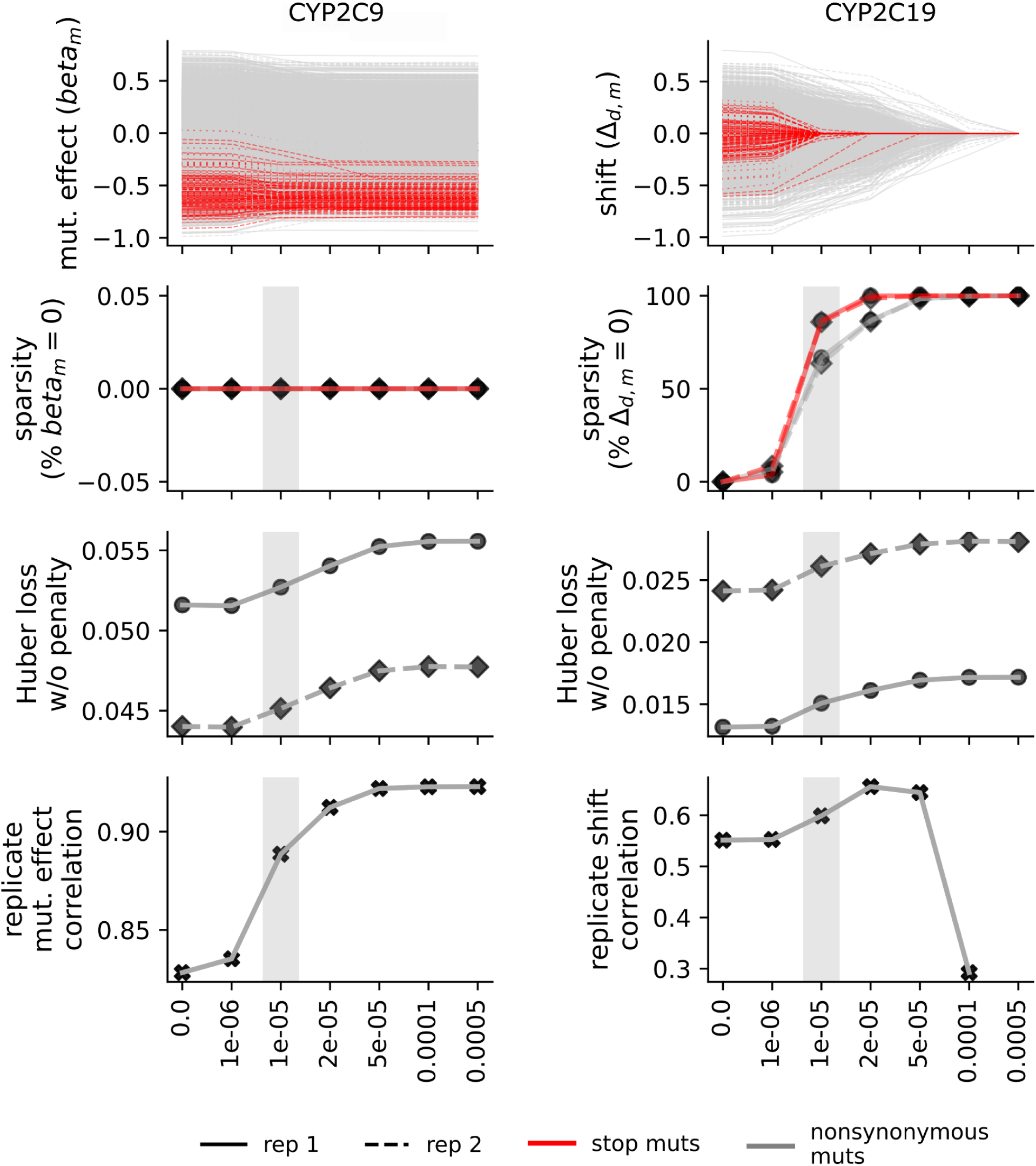
m*u*ltidms shrinkage analysis. Defining characteristics of each independent, replicate model fit compared across seven increasing lasso regularization weights (λ) denoted on the x-axis. The first row shows both (left) the inferred mutation effect values (β) in the CYP2C9 reference experiments, and (right) the corresponding set of mutational effect shifts (Δ) in the CYP2C19 experimental background. Mutations to stop codons are highlighted in red. The second row shows sparsity for the two respective parameter sets (separated by mutation type) as a percentage of values equal to zero. In the third row, we show each model fit’s Huber loss on the training set (before application of the penalty terms in the model objective function). Finally, the last row shows correlation of parameter values between the two replicate models trained on independent datasets.

**Supplemental Figure 7.**
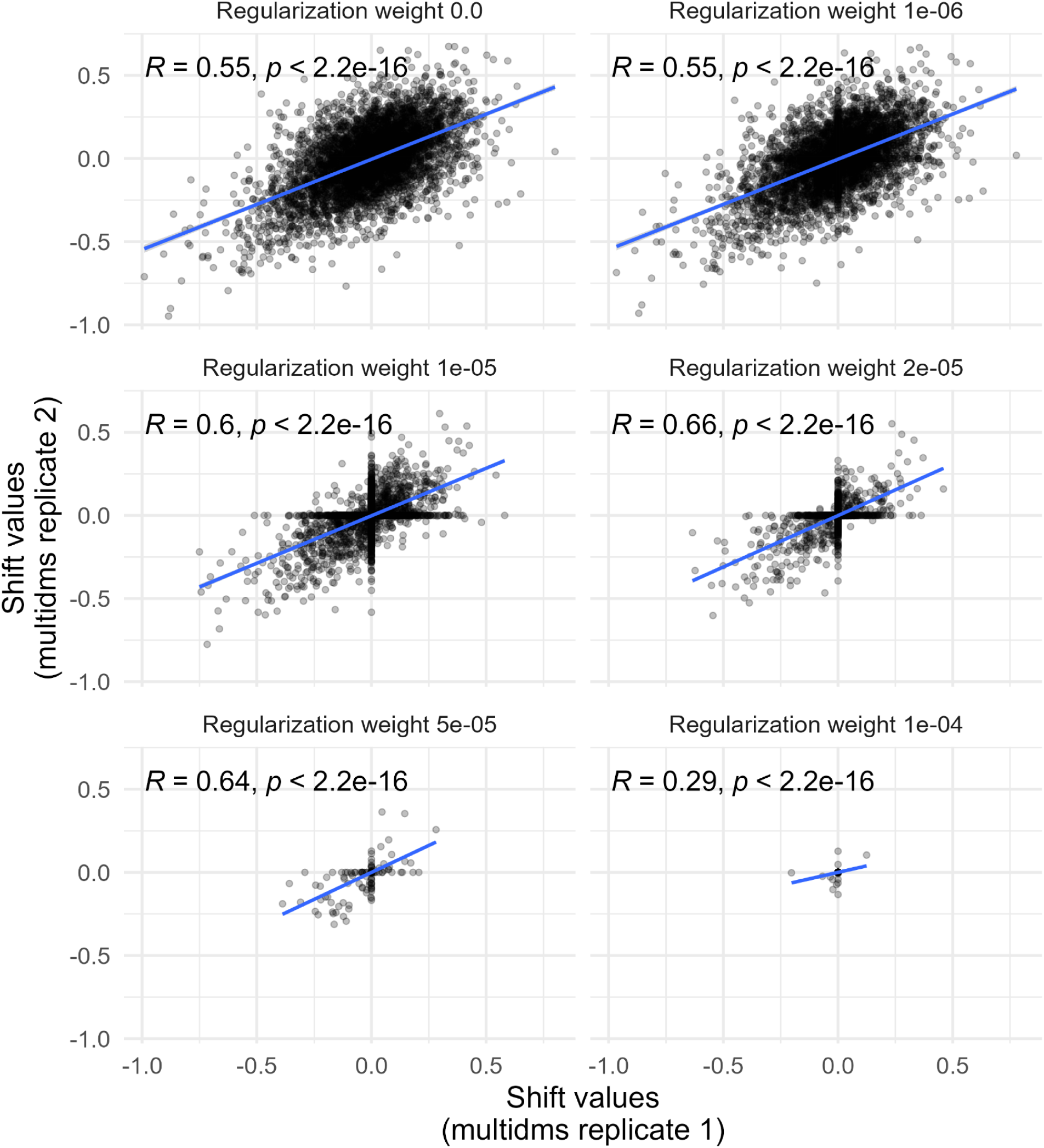
Correlation of *multidms* replicate CYP2C19 shift values. Scatter plot of shift values from two replicate models trained on independent datasets. Regularization weight used during fitting ranges from 0.0 to 1e-04 (left to right, top to bottom). Pearson’s R shown in top left corner of each plot.

**Supplemental Figure 8.**
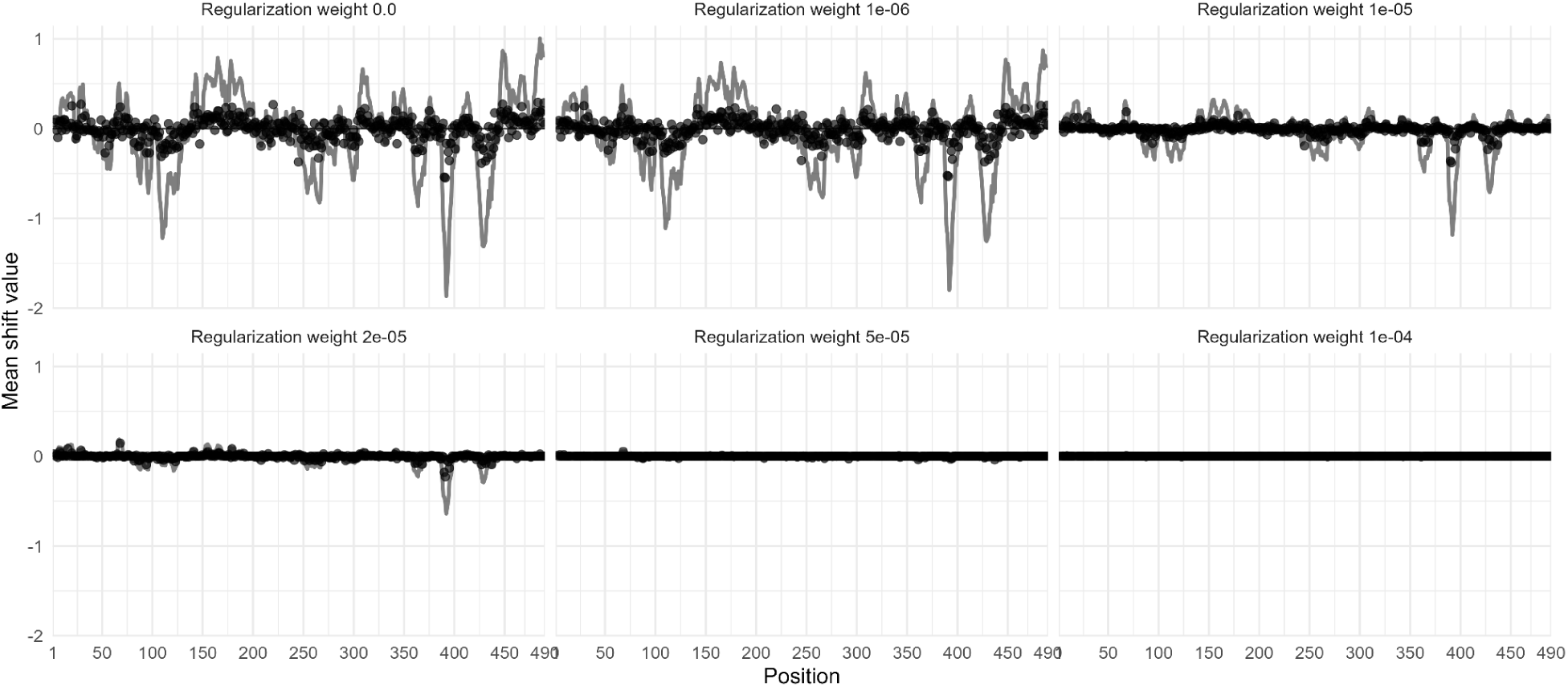
Position means of *multidms* inferred CYP2C19 shift values. Each figure shows individual shifts (Δ) in mutational effect on the CYP2C19 background (relative to the effect in CYP2C9) as a function of mutation site. Gray line is the rolling sum of the mean shift values with k=5 window size. Regularization weight used during fitting ranges from 0.0 to 1e-4 (left to right, top to bottom).

**Supplemental Figure 9.**
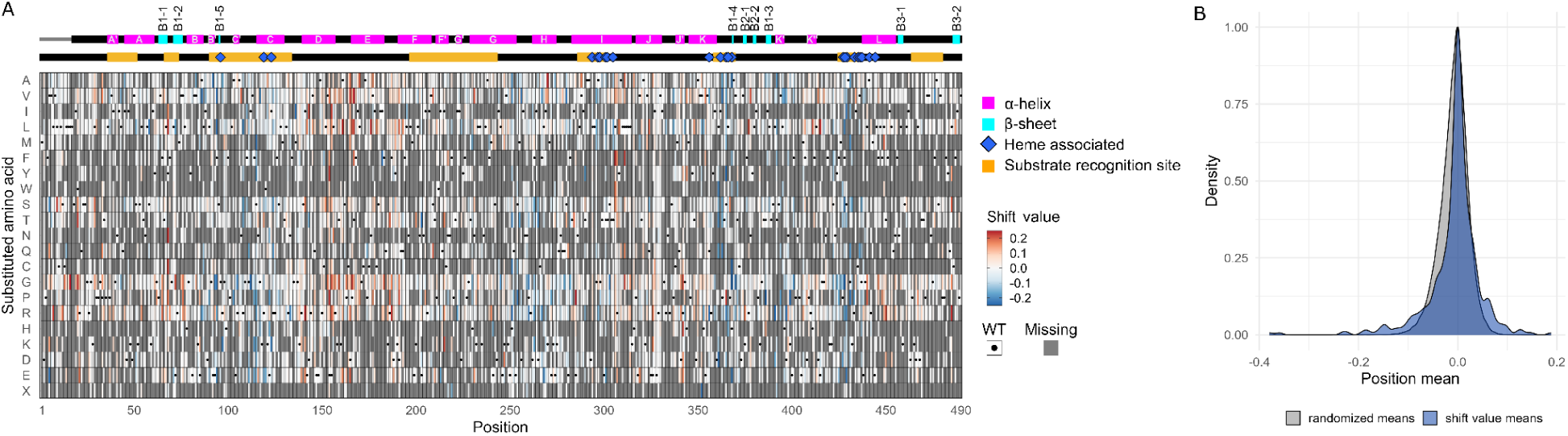
m*u*ltidms shift scores for all variants. A) Heatmap of CYP2C19 shift values across all positions and substituted amino acids. CYP2C19 WT is shown in white with a black dot, and missing data is gray. Substrate recognition regions are shown above the heatmap in orange, and sites that interact with heme are shown with blue diamonds. Secondary structure of CYP2C19 above the heatmap is represented with α-helices shown in magenta and β-sheets shown in cyan. B) Density distribution of mean shift values in blue and randomized mean shift values in gray, as described in Methods.

**Supplemental Figure 10.**
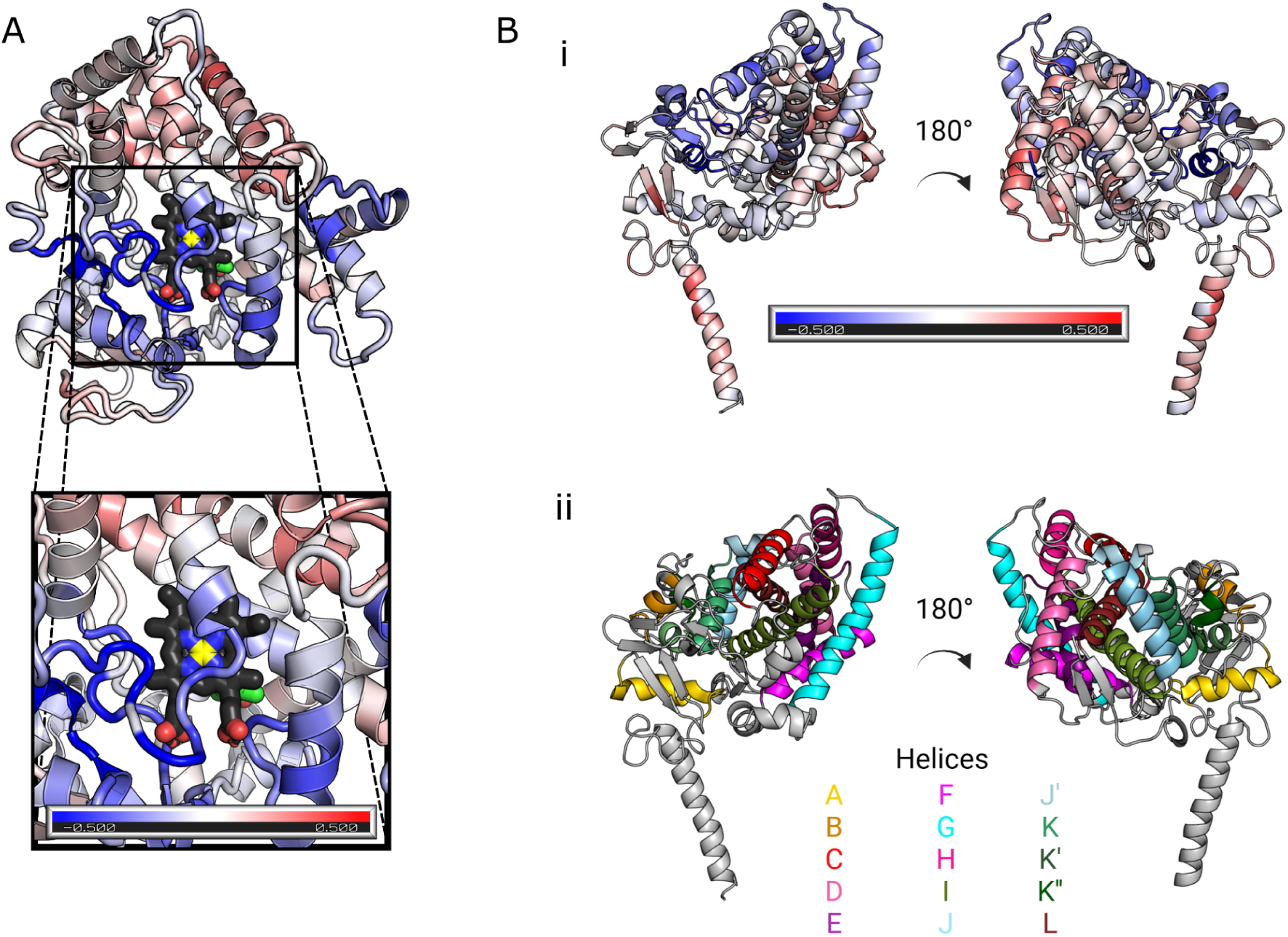
m*u*ltidms shift values plotted on CYP2C19. A) CYP2C19 structure (PDB: <u>4GQS</u>) colored by the rolling sum of position mean shift values (tiled window size k=5, color gradient –0.5 to 0.5). Heme is colored by element (carbon: black, nitrogen: blue, oxygen: red, iron: yellow), and PDB chemical 0XV in green. B) CYP2C19 structure from molecular dynamics simulation ^74^ colored by i) rolling sum of position mean shift values (tiled window size k=5, color gradient –0.5 to 0.5) or ii) α-helix.

**Supplemental Figure 11.**
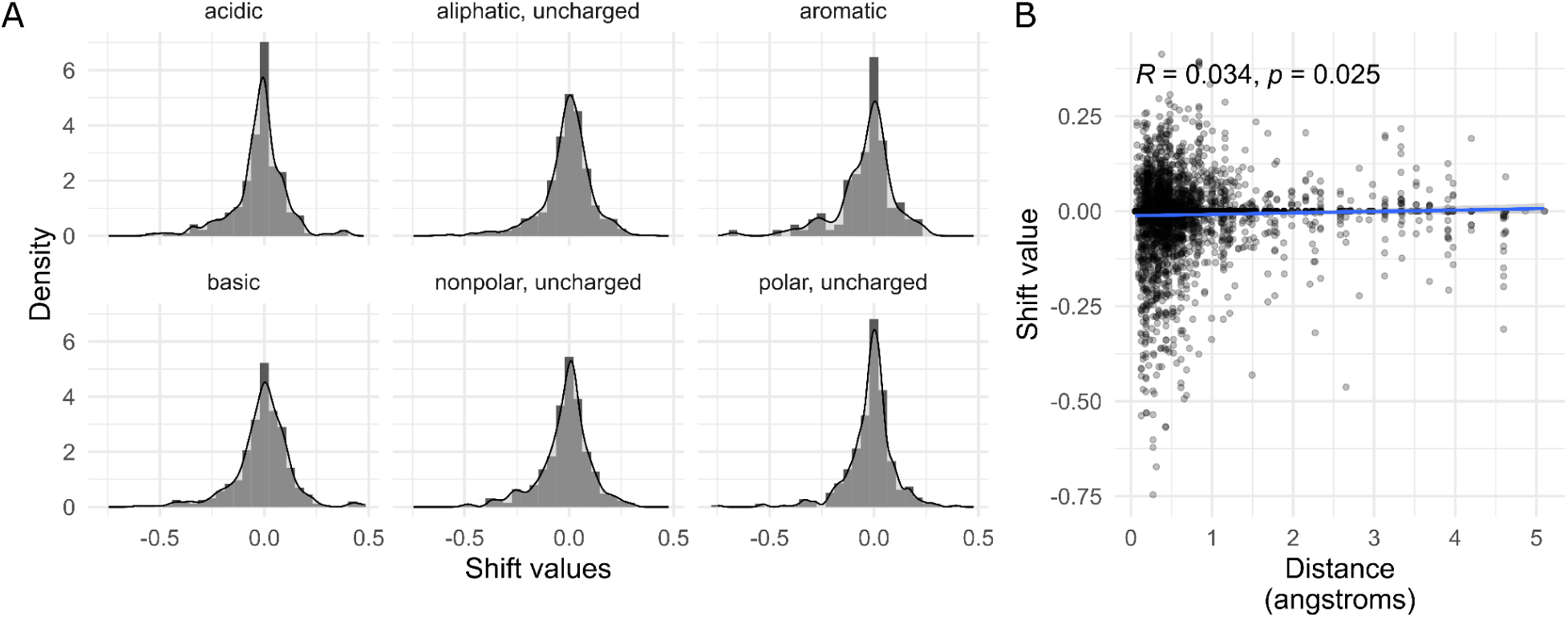
m*u*ltidms shift values across features of CYP2C19 and CYP2C9. A) Density histograms of non-zero shift values across amino acid types. B) Scatter plot showing position mean shift value versus the distance between Cα of aligned CYP2C19 (PDB: <u>4GQS</u>) and CYP2C9 (PDB: <u>1OG2</u>) structures. Pearson’s R = 0.034.

**Supplemental Figure 12.**
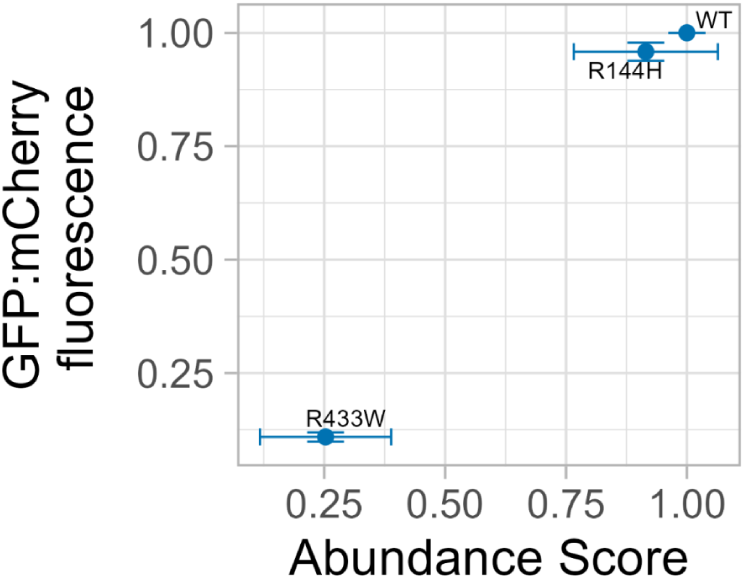
Individual variant validation of *9 (R144H). Geometric mean of the fluorescence signal ratio of GFP:mCherry vs abundance score for CYP2C19 WT, *5 (R433W) with known destabilizing properties, and *9 (R144H). Error bars show standard deviation of fluorescence distributions (y-axis) or standard deviation of abundance scores (x-axis).

**Supplemental Figure 13.**
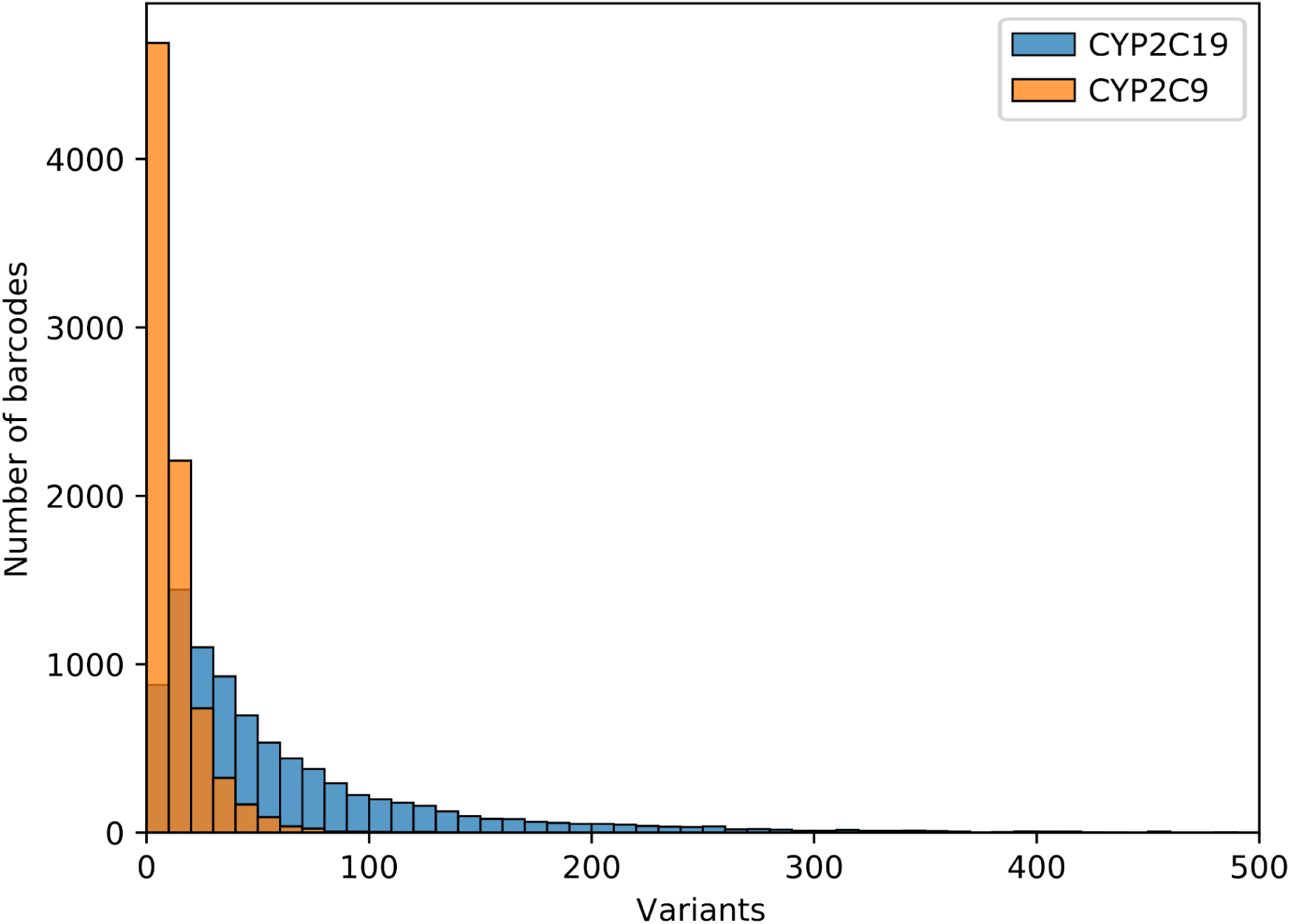
Number of barcodes per variant. Histogram showing the number of unique barcodes associated with individual variants in CYP2C19 (blue) and CYP2C9^30^ (orange) VAMP-seq scans.

## Supporting information

Table S1: CYP2C19 library fluorescence activated cell sorting

Table S2: Library statistics from barcode-variant mapping

Table S3: Plasmids and oligos used in this study

Table S4: CYP2C19 variant abundance scores

Table S5: CYP2C19 star allele CPIC functional annotations

Table S6: CYP2C19 individual variant validation of abundance scores

Table S7: multidms analysis of CYP2C9 and CYP2C19 abundance scores

## Notes

https://github.com/FowlerLab/cyp2c19_2c9

### Competing Interest Statement

The authors have declared no competing interest.

## References

1. Nelson, D. R. Progress in tracing the evolutionary paths of cytochrome P450. Biochim. Biophys. Acta 1814, 14–18 (2011).

2. Munro, A. W., Girvan, H. M., Mason, A. E., Dunford, A. J. & McLean, K. J. What makes a P450 tick? Trends Biochem. Sci. 38, 140–150 (2013).

3. Coon, M. J. Cytochrome P450: nature’s most versatile biological catalyst. Annu. Rev. Pharmacol. Toxicol. 45, 1–25 (2005).

4. Zhao, M. et al. Cytochrome P450 Enzymes and Drug Metabolism in Humans. Int. J. Mol. Sci. 22, (2021).

5. Werck-Reichhart, D. & Feyereisen, R. Cytochromes P450: a success story. Genome Biol. 1, REVIEWS3003 (2000).

6. Sirim, D., Widmann, M., Wagner, F. & Pleiss, J. Prediction and analysis of the modular structure of cytochrome P450 monooxygenases. BMC Struct. Biol. 10, 34 (2010).

7. Reynald, R. L., Sansen, S., Stout, C. D. & Johnson, E. F. Structural characterization of human cytochrome P450 2C19: active site differences between P450s 2C8, 2C9, and 2C19. J. Biol. Chem. 287, 44581–44591 (2012).

8. Niwa, T. & Yamazaki, H. Comparison of cytochrome P450 2C subfamily members in terms of drug oxidation rates and substrate inhibition. Curr. Drug Metab. 13, 1145–1159 (2012).

9. Wishart, D. S. et al. DrugBank 5.0: a major update to the DrugBank database for 2018. Nucleic Acids Res. 46, D1074–D1082 (2018).

10. Mustafa, G., Nandekar, P. P., Bruce, N. J. & Wade, R. C. Differing Membrane Interactions of Two Highly Similar Drug-Metabolizing Cytochrome P450 Isoforms: CYP 2C9 and CYP 2C19. Int. J. Mol. Sci. 20, (2019).

11. Thomson, R. Structural and functional characterisation of ancestral cytochromes P450 from family 2 in tetrapods. (espace.library.uq.edu.au, 2021). doi:10.14264/a159633.

12. Zanger, U. M. & Schwab, M. Cytochrome P450 enzymes in drug metabolism: regulation of gene expression, enzyme activities, and impact of genetic variation. Pharmacol. Ther. 138, 103–141 (2013).

13. Lazarou, J., Pomeranz, B. H. & Corey, P. N. Incidence of adverse drug reactions in hospitalized patients: a meta-analysis of prospective studies. JAMA 279, 1200–1205 (1998).

14. de Vries, E. N., Ramrattan, M. A., Smorenburg, S. M., Gouma, D. J. & Boermeester, M. A. The incidence and nature of in-hospital adverse events: a systematic review. Qual. Saf. Health Care 17, 216–223 (2008).

15. Sultana, J., Cutroneo, P. & Trifirò, G. Clinical and economic burden of adverse drug reactions. J. Pharmacol. Pharmacother. 4, S73–7 (2013).

16. Relling, M. V. & Klein, T. E. CPIC: Clinical Pharmacogenetics Implementation Consortium of the Pharmacogenomics Research Network. Clin. Pharmacol. Ther. 89, 464–467 (2011).

17. Sim, S. C. & Ingelman-Sundberg, M. The Human Cytochrome P450 (CYP) Allele Nomenclature website: a peer-reviewed database of CYP variants and their associated effects. Hum. Genomics 4, 278–281 (2010).

18. Klein, M. D., Lee, C. R. & Stouffer, G. A. Clinical outcomes of CYP2C19 genotype-guided antiplatelet therapy: existing evidence and future directions. Pharmacogenomics 19, 1039–1046 (2018).

19. Galli, M. et al. Guided versus standard antiplatelet therapy in patients undergoing percutaneous coronary intervention: a systematic review and meta-analysis. Lancet 397, 1470–1483 (2021).

20. Pereira, N. L. et al. Effect of CYP2C19 Genotype on Ischemic Outcomes During Oral P2Y12 Inhibitor Therapy: A Meta-Analysis. JACC Cardiovasc. Interv. 14, 739–750 (2021).

21. Dean, L. & Kane, M. Clopidogrel Therapy and CYP2C19 Genotype. (National Center for Biotechnology Information (US), 2022).

22. Matreyek, K. A. et al. Multiplex assessment of protein variant abundance by massively parallel sequencing. Nat. Genet. 50, 874–882 (2018).

23. Yen, H.-C. S., Xu, Q., Chou, D. M., Zhao, Z. & Elledge, S. J. Global protein stability profiling in mammalian cells. Science 322, 918–923 (2008).

24. Zutz, A. et al. A dual-reporter system for investigating and optimizing protein translation and folding in E. coli. Nat. Commun. 12, 6093 (2021).

25. Kim, I., Miller, C. R., Young, D. L. & Fields, S. High-throughput analysis of in vivo protein stability. Mol. Cell. Proteomics 12, 3370–3378 (2013).

26. Klesmith, J. R., Bacik, J.-P., Wrenbeck, E. E., Michalczyk, R. & Whitehead, T. A. Trade-offs between enzyme fitness and solubility illuminated by deep mutational scanning. Proc. Natl. Acad. Sci. U. S. A. 114, 2265–2270 (2017).

27. Hargrove, J. L. & Schmidt, F. H. The role of mRNA and protein stability in gene expression. FASEB J. 3, 2360–2370 (1989).

28. Christensen, S., Wernersson, C. & André, I. Facile Method for High-throughput Identification of Stabilizing Mutations. J. Mol. Biol. 435, 168209 (2023).

29. Suiter, C. C. et al. Massively parallel variant characterization identifies *NUDT15* alleles associated with thiopurine toxicity. Proceedings of the National Academy of Sciences 117, 5394–5401 (2020).

30. Amorosi, C. J. et al. Massively parallel characterization of CYP2C9 variant enzyme activity and abundance. Am. J. Hum. Genet. 108, 1735–1751 (2021).

31. DePristo, M. A., Weinreich, D. M. & Hartl, D. L. Missense meanderings in sequence space: a biophysical view of protein evolution. Nat. Rev. Genet. 6, 678–687 (2005).

32. Matreyek, K. A., Stephany, J. J. & Fowler, D. M. A platform for functional assessment of large variant libraries in mammalian cells. Nucleic Acids Res. 45, e102 (2017).

33. Matreyek, K. A., Stephany, J. J., Chiasson, M. A., Hasle, N. & Fowler, D. M. An improved platform for functional assessment of large protein libraries in mammalian cells. Nucleic Acids Res. 48, e1 (2020).

34. Zhang, L. et al. CYP2C9 and CYP2C19: Deep Mutational Scanning and Functional Characterization of Genomic Missense Variants. Clin. Transl. Sci. 13, 727–742 (2020).

35. Hasemann, C. A., Kurumbail, R. G., Boddupalli, S. S., Peterson, J. A. & Deisenhofer, J. Structure and function of cytochromes P450: a comparative analysis of three crystal structures. Structure 3, 41–62 (1995).

36. Mestres, J. Structure conservation in cytochromes P450. Proteins 58, 596–609 (2005).

37. Gricman, Ł., Vogel, C. & Pleiss, J. Identification of universal selectivity-determining positions in cytochrome P450 monooxygenases by systematic sequence-based literature mining. Proteins 83, 1593–1603 (2015).

38. Gricman, Ł., Vogel, C. & Pleiss, J. Conservation analysis of class-specific positions in cytochrome P450 monooxygenases: functional and structural relevance. Proteins 82, 491–504 (2014).

39. Berka, K., Hendrychová, T., Anzenbacher, P. & Otyepka, M. Membrane position of ibuprofen agrees with suggested access path entrance to cytochrome P450 2C9 active site. J. Phys. Chem. A 115, 11248–11255 (2011).

40. Lertkiatmongkol, P. et al. Distal effect of amino acid substitutions in CYP2C9 polymorphic variants causes differences in interatomic interactions against (S)-warfarin. PLoS One 8, e74053 (2013).

41. Foti, R. S. et al. Ligand-based design of a potent and selective inhibitor of cytochrome P450 2C19. J. Med. Chem. 55, 1205–1214 (2012).

42. Altarsha, M., Benighaus, T., Kumar, D. & Thiel, W. How is the reactivity of cytochrome P450cam affected by Thr252X mutation? A QM/MM study for X = serine, valine, alanine, glycine. J. Am. Chem. Soc. 131, 4755–4763 (2009).

43. Haines, D. C., Tomchick, D. R., Machius, M. & Peterson, J. A. Pivotal role of water in the mechanism of P450BM-3. Biochemistry 40, 13456–13465 (2001).

44. Nair, P. C., McKinnon, R. A. & Miners, J. O. Cytochrome P450 structure-function: insights from molecular dynamics simulations. Drug Metab. Rev. 48, 434–452 (2016).

45. Niwa, T. et al. Amino acid residues affecting the activities of human cytochrome P450 2C9 and 2C19. Drug Metab. Dispos. 30, 931–936 (2002).

46. Goldstein, J. A. & de Morais, S. M. Biochemistry and molecular biology of the human CYP2C subfamily. Pharmacogenetics 4, 285–299 (1994).

47. Gumulya, Y. et al. Engineering highly functional thermostable proteins using ancestral sequence reconstruction. Nature Catalysis 1, 878–888 (2018).

48. Haddox, H. K. et al. Jointly modeling deep mutational scans identifies shifted mutational effects among SARS-CoV-2 spike homologs. bioRxiv 2023.07.31.551037 (2023) doi:10.1101/2023.07.31.551037.

49. Denisov, I. G., Makris, T. M., Sligar, S. G. & Schlichting, I. Structure and chemistry of cytochrome P450. Chem. Rev. 105, 2253–2277 (2005).

50. Gotoh, O. Substrate recognition sites in cytochrome P450 family 2 (CYP2) proteins inferred from comparative analyses of amino acid and coding nucleotide sequences. J. Biol. Chem. 267, 83–90 (1992).

51. Jung, F. et al. Identification of amino acid substitutions that confer a high affinity for sulfaphenazole binding and a high catalytic efficiency for warfarin metabolism to P450 2C19. Biochemistry 37, 16270–16279 (1998).

52. Attia, T. Z. et al. Effect of cytochrome P450 2C19 and 2C9 amino acid residues 72 and 241 on metabolism of tricyclic antidepressant drugs. Chem. Pharm. Bull. 62, 176–181 (2014).

53. Wada, Y. et al. Important amino acid residues that confer CYP2C19 selective activity to CYP2C9. J. Biochem. 144, 323–333 (2008).

54. Klose, T. S. et al. Identification of residues 286 and 289 as critical for conferring substrate specificity of human CYP2C9 for diclofenac and ibuprofen. Arch. Biochem. Biophys. 357, 240–248 (1998).

55. Ibeanu, G. C. et al. Identification of residues 99, 220, and 221 of human cytochrome P450 2C19 as key determinants of omeprazole activity. J. Biol. Chem. 271, 12496–12501 (1996).

56. Lewis, D. F. et al. Molecular modelling of human CYP2C subfamily enzymes CYP2C9 and CYP2C19: rationalization of substrate specificity and site-directed mutagenesis experiments in the CYP2C subfamily. Xenobiotica 28, 235–268 (1998).

57. Tsao, C. C. et al. Identification of human CYP2C19 residues that confer S-mephenytoin 4’-hydroxylation activity to CYP2C9. Biochemistry 40, 1937–1944 (2001).

58. Goulding, R., Dawes, D., Price, M., Wilkie, S. & Dawes, M. Genotype-guided drug prescribing: a systematic review and meta-analysis of randomized control trials. Br. J. Clin. Pharmacol. 80, 868–877 (2015).

59. Schmiedl, S. et al. Preventable ADRs leading to hospitalization – results of a long-term prospective safety study with 6,427 ADR cases focusing on elderly patients. Expert Opin. Drug Saf. 17, 125–137 (2018).

60. Ibeanu, G. C. et al. Identification of new human CYP2C19 alleles (CYP2C19*6 and CYP2C19*2B) in a Caucasian poor metabolizer of mephenytoin. J. Pharmacol. Exp. Ther. 286, 1490–1495 (1998).

61. Derayea, S. M. et al. Impact of single nucleotide polymorphisms (R132Q and W120R) on the binding affinity and metabolic activity of CYP2C19 toward some therapeutically important substrates. Xenobiotica 50, 1510–1519 (2020).

62. Blaisdell, J. et al. Identification and functional characterization of new potentially defective alleles of human CYP2C19. Pharmacogenetics 12, 703–711 (2002).

63. Fayer, S. et al. Closing the gap: Systematic integration of multiplexed functional data resolves variants of uncertain significance in BRCA1, TP53, and PTEN. Am. J. Hum. Genet. 108, 2248–2258 (2021).

64. Ionova, Y. et al. CYP2C19 Allele Frequencies in Over 2.2 Million Direct-to-Consumer Genetics Research Participants and the Potential Implication for Prescriptions in a Large Health System. Clin. Transl. Sci. 13, 1298–1306 (2020).

65. Peng, C.-C., Cape, J. L., Rushmore, T., Crouch, G. J. & Jones, J. P. Cytochrome P450 2C9 type II binding studies on quinoline-4-carboxamide analogues. J. Med. Chem. 51, 8000–8011 (2008).

66. Polgár, T., Menyhárd, D. K. & Keseru, G. M. Effective virtual screening protocol for CYP2C9 ligands using a screening site constructed from flurbiprofen and S-warfarin pockets. J. Comput. Aided Mol. Des. 21, 539–548 (2007).

67. Skopalík, J., Anzenbacher, P. & Otyepka, M. Flexibility of human cytochromes P450: molecular dynamics reveals differences between CYPs 3A4, 2C9, and 2A6, which correlate with their substrate preferences. J. Phys. Chem. B 112, 8165–8173 (2008).

68. Jain, P. C. & Varadarajan, R. A rapid, efficient, and economical inverse polymerase chain reaction-based method for generating a site saturation mutant library. Anal. Biochem. 449, 90–98 (2014).

69. Zhang, J., Kobert, K., Flouri, T. & Stamatakis, A. PEAR: a fast and accurate Illumina Paired-End reAd mergeR. Bioinformatics 30, 614–620 (2014).

70. Zhao, L., Liu, Z., Levy, S. F. & Wu, S. Bartender: a fast and accurate clustering algorithm to count barcode reads. Bioinformatics 34, 739–747 (2018).

71. Yeh, C.-L. C., Amorosi, C. J., Showman, S. & Dunham, M. J. PacRAT: a program to improve barcode-variant mapping from PacBio long reads using multiple sequence alignment. Bioinformatics 38, 2927–2929 (2022).

72. García-Nafría, J., Watson, J. F. & Greger, I. H. IVA cloning: A single-tube universal cloning system exploiting bacterial In Vivo Assembly. Sci. Rep. 6, 27459 (2016).

73. Benjamini, Y. & Hochberg, Y. Controlling the False Discovery Rate: A Practical and Powerful Approach to Multiple Testing. J. R. Stat. Soc. Series B Stat. Methodol. 57, 289–300 (1995).

74. Mustafa, G., Nandekar, P. P., Mukherjee, G., Bruce, N. J. & Wade, R. C. The Effect of Force-Field Parameters on Cytochrome P450-Membrane Interactions: Structure and Dynamics. Sci. Rep. 10, 7284 (2020).

